# NSUN2 drives intestinal stem cell expansion and colorectal tumour initiation via MAPK/ERK signalling

**DOI:** 10.1101/2025.04.11.648351

**Authors:** Aslıhan Bastem Akan, Caroline V. Billard, Szu-Ying Chen, Po-Hsien Huang, Patrizia Cammareri, Adam E. Hall, Paula Preyzner, Farhat Din, Kevin B. Myant

## Abstract

Colorectal cancer is initiated by loss of the *APC* gene, which drives expansion of LGR5+ intestinal stem cell (ISC) populations. Whilst LGR5+ ISC expansion is a critical step for tumour initiation and progression, its regulation is poorly understood. Emerging evidence suggests post-transcriptional RNA modifications play a key role in cancer biology, but their role in CRC initiation has not been explored. Here, we identify the m^5^C methyltransferase NSUN2 as a key regulator of ISC expansion and intestinal tumourigenesis. NSUN2 is upregulated in multiple CRC mouse models and human tumours and its depletion impairs ISC expansion and hyperproliferation, leading to reduced tumour initiation. Transcriptome-wide bisulphite sequencing revealed that NSUN2-mediates m^5^C methylation on mRNAs encoding key ISC regulators and components of the MAPK/ERK pathway. Mechanistically, loss of NSUN2 reduces ERK phosphorylation in *Apc-*deficient models and oncogenic *Kras^G12D^*expression is sufficient to restore ERK signalling and rescue ISC expansion. Together, this establishes NSUN2 as a key regulator of ISC-driven CRC initiation and describes a novel molecular mechanism linking m^5^C methylation to MAPK driven stem cell transformation.

## Introduction

Colorectal cancer (CRC) is the third commonest cancer worldwide, with nearly two million new cases annually, and the second commonest cause of cancer-related death^1^. CRC initiation is driven by malignant transformation of LGR5+ intestinal stem cells (ISCs), which maintain normal intestinal homeostasis through constant self-renewal and differentiation^2,3^. This process is normally tightly controlled by signalling pathways such as WNT, MAPK, BMP and TGF-β. However, during tumourigenesis these pathways become dysregulated leading to ISC expansion and tumour formation^4^. Most CRCs are initiated by loss of the *APC* gene, leading to a hyper-proliferative early stage, and progress to metastatic carcinoma via accumulation of additional mutations in genes including *TP53, KRAS* and *SMAD4*^5–9^. Although targeted therapies and immunotherapies have improved outcomes for some patients, CRC remains difficult to treat due to therapy resistance and disease relapse^10–12^. This is often driven by the activity of cancer stem cells, a subpopulation of tumour-initiating cells which arise from transformed ISCs via aberrant genetic or epigenetic pathways and which sustain tumour growth and evade conventional treatments^13–16^.

Post translational modifications of RNA are important regulators of tumourigenesis. These modifications are found in distinct RNA types and influence a multitude of diverse functions, such as gene expression, RNA stability, RNA localisation and protein translation ^17,18^ ^18–20^ ^21,22^. Over 170 RNA modifications that control fate of RNA and mediate cellular functions have been identified^23^ . Of these, 5-methylcytidine (m^5^C) is one of the best described and was first discovered by Amos and Korn^24^. m^5^C is widely found in tRNAs and rRNAs, where it stabilises RNA structure and regulates translation efficiency. More recently m5C has been identified in mRNA, where it modulates RNA stability^25–27^, export^28^ and translation^18^. RNA modifications are regulated by diverse proteins categorised as writers, readers and erasers^29^. m^5^C can be introduced to RNAs by the NSUN family of proteins (NSUN1 to NSUN7) or DNMT2^30^ and NSUN2 and NSUN6 have been shown to modulate mRNA methylation^30–32^. Of the NSUN proteins, NSUN2 has gained increasing attention due to its oncogenic potential.

NSUN2 was first identified as a target of the well-known proto-oncogene MYC in the epidermis^33,34^. It catalyses m^5^C methylation in tRNAs, rRNAs and, described more recently, mRNAs^25,35–44^. NSUN2- mediated m^5^C has been shown to regulate RNA stability, translation efficiency and cellular differentiation. Multiple research groups have reported increased NSUN2 mRNA and protein expression in various cancers, including gastric, oesophageal, hepatocellular, bladder, osteosarcoma, and cervical cancers. Additionally, NSUN2 has been demonstrated to play a role in regulating tumour growth, migration, and invasion^26,45–49^. However, the role of NSUN2, and that of mRNA m^5^C methylation, in intestinal homeostasis and colorectal tumourigenesis remains poorly understood. Therefore, a better understanding of the role of m^5^C mRNA methylation and its role in ISC expansion and tumour initiation is crucial for fully defining the mechanisms mediating tumour initiation.

In this study we define the role of NSUN2 in CRC initiation using *Apc*-deficient CRC mouse models. We find that NSUN2 is overexpressed following *Apc-*loss and in human CRC and correlates with poor disease outcome. Functionally, we find that NSUN2 depletion impairs ISC-driven hyperproliferation and cancer stemness in *Apc*-deficient *in vitro* organoids and *in vivo* cancer models, without affecting normal intestinal homeostasis. Transcriptome-wide bisulphite analysis reveals that loss of NSUN2 leads to bidirectional changes in m^5^C mRNA methylation, particularly in mRNAs encoding ISC regulators and MAPK/ERK components. Furthermore, we find that MAPK/ERK pathway activation in *Apc-*deficient organoids depends on NSUN2 expression, an effect rescued by expression of oncogenic *Kras^G12D^*. Together, these findings establish NSUN2-mediated m^5^C methylation as a crucial regulator of ISC expansion and CRC initiation, for the first-time linking RNA modifications to colorectal stem cell transformation.

## Results

### NSUN2, 5-methylcytidine writer, is upregulated in *Apc-loss-driven* hyper-proliferative intestine

To better understand the role of m^5^C methylation in Wnt-driven tumour initiation we analysed our previously generated RNA-seq dataset derived from intestinal epithelium of *Vil-creERT2 Apc^fl/fl^*mice following tamoxifen-induced *Apc* deletion^50^. This analysis revealed significant overexpression of the m^5^C methyltransferase *Nsun2,* but not other known RNA m^5^C methyltransferases, readers or eraser enzymes, following *Apc* loss (Supplementary Fig S1A). To further characterise NSUN2 expression in colorectal tumourigenesis, we performed IHC staining (Fig 1A-1B) and WB (Fig 1C-1D and Supplementary Fig S1B-C) for NSUN2 in normal and *Apc*-deficient mouse intestines which indicated elevated NSUN2 expression in hyper-proliferative intestines. We next investigated NSUN2 expression in intestinal adenomas and found increased NSUN2 expression in adenomas arising in the colon from the spontaneous *VilCreERT2-Apc^+/-^* model (Fig 1E-1F)^51^. To analyse NSUN2 expression following malignant stem cell transformation, we used the tamoxifen-inducible *Lgr5-IRES-eGFP-CreERT2 Apc^fl/fl^* model where both copies of *Apc* are deleted in *Lgr5*+ stem cells^4^. Again, NSUN2 expression was elevated in adenomas arising from Wnt hyperactivation in stem cells (Fig 1G-H). Together, these data demonstrate elevated expression of NSUN2 in *Apc-*deficient CRC models, suggesting a potential role in tumourigenesis.

**Figure 1.**
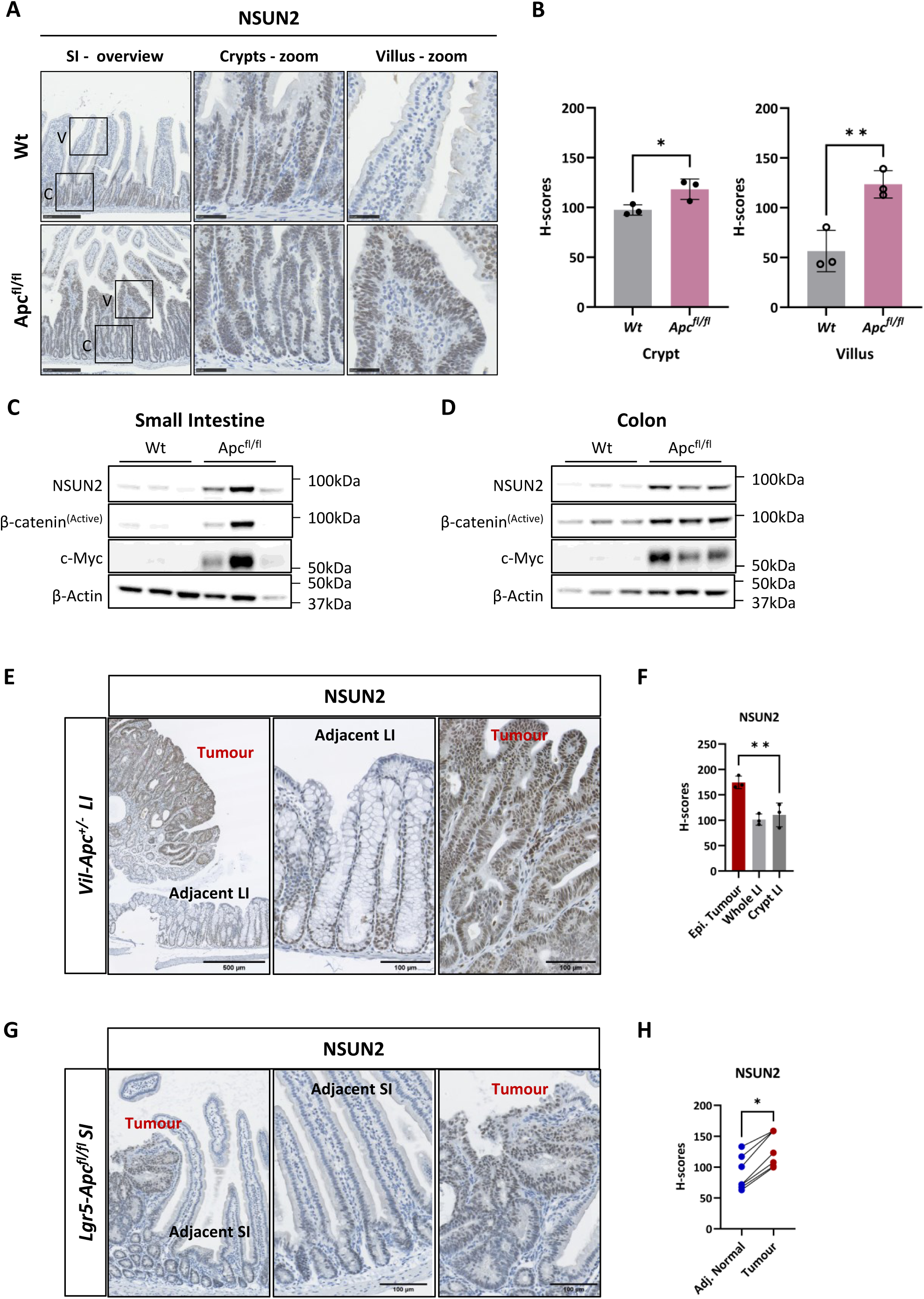
RNA 5-methylcytidine writer, NSUN2, is upregulated following *Apc-loss in* mouse small and large intestinal tissues. **A** Representative images of NSUN2 expression using immunohistochemistry method in wild-type and Apc^fl/fl^ small intestinal tissues. C represents crypt and V represents villus. Scale bars are shown as 500 µm (overview) and 100 µm (magnified). **B** Histo-score quantification of NSUN2 expression using immunohistochemistry method in wild-type and Apc^fl/fl^ small intestinal tissues (data are presented as mean ± SD; *p:0.0356, **p:0.0096; two-tailed t-test, n = 3vs3 biologically independent mice). **C** Images of western blotting analysis for NSUN2, Active β-Catenin, c-Myc and a house-keeping control β-ACTIN protein expressions in wild-type and Apc^fl/fl^ mouse small intestines. **D** Images of western blotting analysis for NSUN2, Active β-Catenin, c-Myc and a house-keeping control β-ACTIN protein expressions in wild-type and Apc^fl/fl^ mouse colons. **E** Representative immunohistochemistry images of NSUN2 expression in normal adjacent tissue and tumours tissue from Vil-CreERT2-*Apc*^+/-^ mouse colon **F** Histo-score quantification of (E). Data are presented as mean ± SD; **p:0.0058; Ordinary One-way ANOVA test compared to Epithelial tumour, n = 3 biologically independent mice. **G** Representative immunohistochemistry images of NSUN2 expression in normal adjacent tissue and tumours tissue from Lgr5-IRES-eGFP-CreERT2-Apc^fl/fl^ mouse small intestines. **H** Histo-score quantification of (G). Data are presented as mean ± SD; *p:0.0189; two-tailed t-test, n=7 adjacent normal tissue vs n=7 tumour tissue parts used from n = 3 biologically independent mice.

### NSUN2 is overexpressed in human CRC and high expression of NSUN2 correlates with poor prognosis

Next, we utilised The Cancer Genome Atlas Colon (TCGA-COAD) and Rectum Adenocarcinoma (TCGA-READ) datasets in order to investigate the expression of NSUN2 in colorectal cancer patients. In TCGA-COAD patients, NSUN2 is higher in primary colon tumour samples compared to normal colon (Fig 2A; p<0.0001) and correlates with poor overall (Fig 2B; p=0.035) and relapse-free survival (Fig 2C; p=0.007). Similar NSUN2 expression and survival correlation was observed in TCGA-READ patients, except for overall survival (Fig 2D-F). Given that *APC, KRAS* and *P53* are key drivers in the ‘adenoma-to-carcinoma’ sequence model of CRC, we next analysed *NSUN2* expression in relation to these mutations^6^. Therefore, we next analysed *NSUN2* expression in patients of TCGA-COAD and TCGA-READ datasets in case of mutant and wild-type version of these major driver genes. The results showed that NSUN2 is overexpressed in patients bearing *APC*-truncation (left, p=0.0017, n=139 vs 385) and *KRAS*-mutation (middle, p=0.0011, n=308 vs 212), but is not significantly associated with *P53*-mutation (right, p=0.0717, n=211 vs 309) (Supplementary Fig S2A). Altogether, these findings indicate that *NSUN2* expression is increased in colorectal cancer and correlates with poor prognosis.

**Figure 2.**
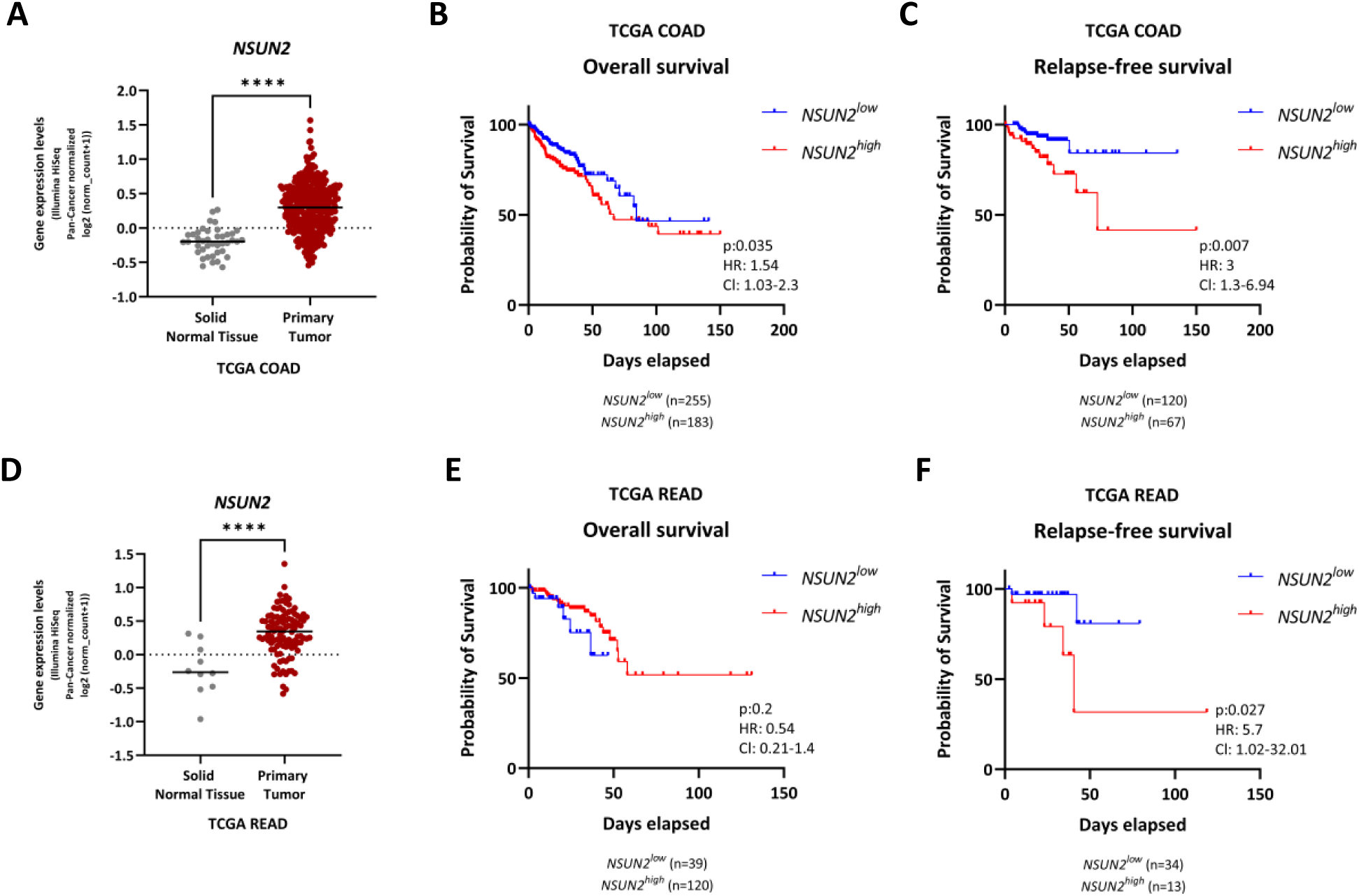
NSUN2 is overexpressed in patient primary tumour compared to adjacent normal tissue and high NSUN2 correlated with poor survival. **A** *NSUN2* mRNA expression is detected in primary tumours of TCGA COAD patients compared to normal adjacent tissues (NAT) in Xenabrowser platform^69^. log2(normalised_count+1) gene expression levels are represented by Illumina Hiseq. z-scores represent normalised gene expression in tumour samples over a reference (normal) samples (data are presented as mean ± SD; ****p<0.0001; two-tailed t-test, n=41 normal adjacent tissue vs n=325 primary tumour). **B-C** Overall survival and Relapse-free survival of patients in TCGA COAD dataset based on NSUN2 expression. Overall and Relapse-free survival of human CRC patients separated by high vs low/intermediate *NSUN2* expression. The red line represents *NSUN2^high^*, and the blue line represents *NSUN2^low^* patients. Log-rank P value, hazard ratio (HR) and 95% confidence intervals (CI) presented. D *NSUN2* mRNA expression is detected in primary tumours of TCGA READ patients compared to normal adjacent tissues (NAT) in Xenabrowser platform^69^. log2(normalised_count+1) gene expression levels are represented by Illumina Hiseq. z-scores represent normalised gene expression in tumour samples over a reference (normal) samples (data are presented as mean ± SD; ****p<0.0001; two-tailed t-test, n=10 normal adjacent tissue vs n=104 primary tumour). **E-F** Overall survival and Relapse-free survival of patients in TCGA READ dataset based on NSUN2 expression. Overall and Relapse-free survival of human CRC patients separated by high vs low/intermediate *NSUN2* expression. The red line represents *NSUN2^high^*, and the blue line represents *NSUN2^low^* patients. Log-rank P value, hazard ratio (HR) and 95% confidence intervals (CI) presented.

### *Nsun2* depletion reduces self-renewal of *Apc^fl/fl^* 3D mouse organoids

To evaluate the phenotypic role of NSUN2, we knocked down gene expression in *Apc^fl/fl^* organoids using shRNA. We transduced *Apc^fl/fl^*organoids with control and two independent sh*Nsun2*-targeting lentiviral particles; hereafter named shCtrl, shNsun2-1 and shNsun2-2 respectively. RNA and protein analysis demonstrated effective knockdown of NSUN2 in this organoid model (Fig 3A-B and Supplementary Figure S3A). Dot blot analysis showed this silencing of *Nsun2* resulted in a functional reduction of global RNA m^5^C methylation in *Apc^fl/fl^* organoids (Fig 3C-D). To determine the phenotypic effects of *Nsun2* and RNA m^5^C methylation depletion on organoid growth we carried out colony formation assays. The ability of *Apc^fl/fl^* single cells to form organoids was significantly decreased following *Nsun2*-depletion (Fig 3E-F). In addition to clonogenic capacity, organoid size and overall cell viability measured by Resazurin assay^52^ was reduced in *Apc^shNsun2^*organoids (Fig 3G-H). Since clonogenic capacity is highly related to self-renewal capacity and stem cell activity, we examined mRNA expression of stem cell marker genes in *Nsun2*-depleted *Apc^fl/fl^* organoids. Several well-characterised intestinal stem cell markers such as *Lgr5, Ascl2, Igfbp4, Lrig1*, and *Smoc2* were significantly decreased following *Nsun2* depletion, indicative of a loss of stem cell capacity (Fig 3I). Thus, *Nsun2* is essential for maintaining cancer stem cell function in a Wnt-driven CRC organoid model, potentially through its role in m^5^C methylation.

**Figure 3.**
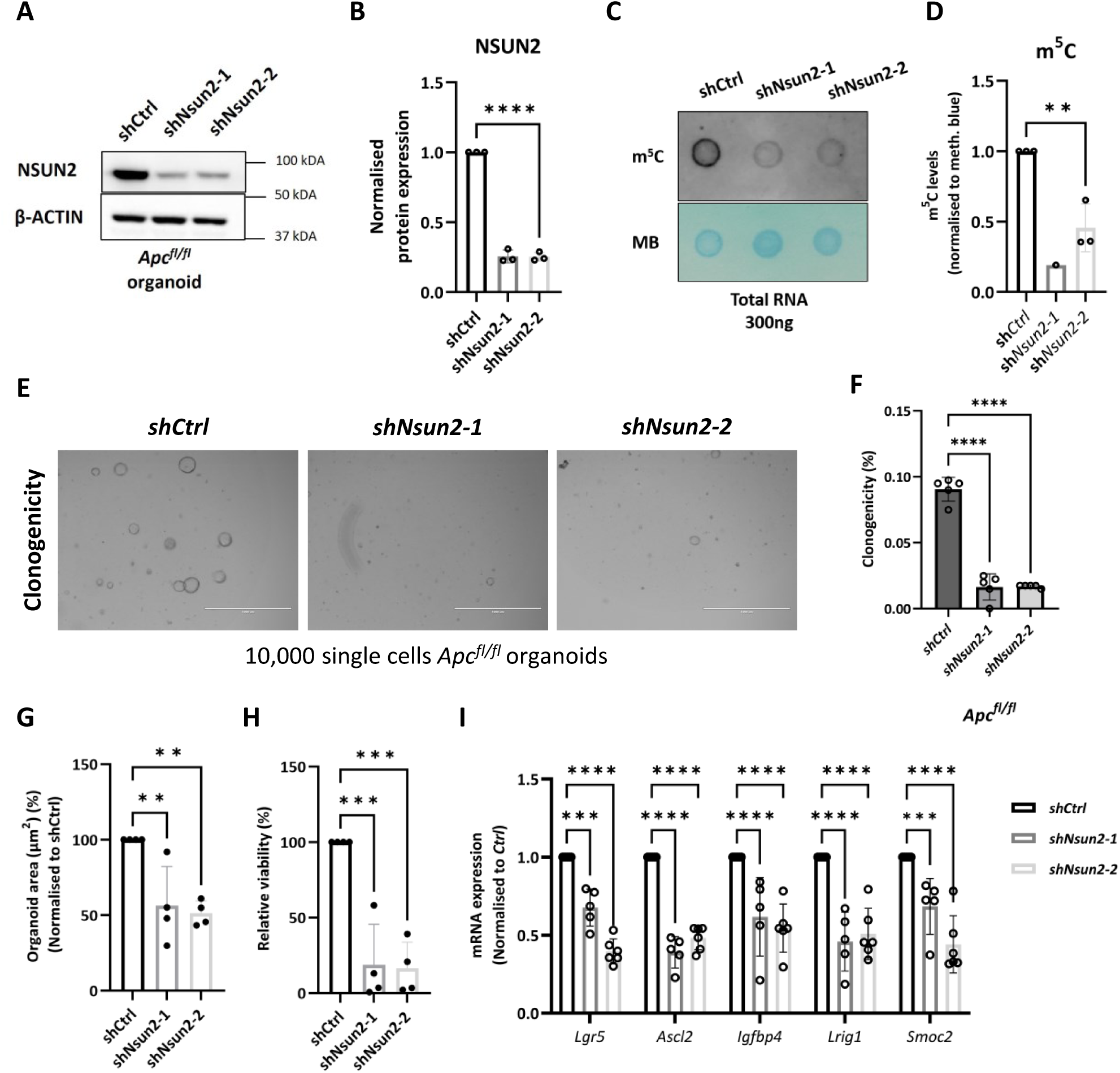
NSUN2 depletion reduces stemness in *Apc^fl/fl^* mouse intestinal organoids. **A** Representative image of validation of Nsun2 knockdown in Vil-CreERT2 mouse *Apc^fl/fl^-shCtrl* and *Apc^fl/fl^-shNsun2* mouse small intestinal organoids by western blot. **B** Quantification of normalised NSUN2 protein expression levels in Vil-CreERT2 mouse *Apc^fl/fl^-shCtrl* and *Apc^fl/fl^-shNsun2* mouse small intestinal organoids by western blot, using β-ACTIN loading control (data are presented as mean ± SD; ****p<0.0001, Ordinary One-way ANOVA test compared to *shCtrl*, n = 3vs3 biologically independent mice). **C** Representative image of 5-methylcytidine (m^5^C) levels in total RNA extracted from *shNsun2-1* and *shNsun2-2* compared to *shCtrl* in *Apc^fl/fl^* mouse small intestinal organoids measured by dot blot assay. Methylene blue used as RNA loading control. **D** The levels of 5-methylcytidine (m^5^C) in total RNA extracted from *shNsun2-1* and *shNsun2-2* compared to *shCtrl* in *Apc^fl/fl^* mouse small intestinal organoids measured by dot blot assay. (data are presented as mean ± SD; **p:0.0092, Ordinary One-way ANOVA test compared to *shCtrl*, n = 3vs1vs3 biologically independent mice). **E** Representative image of Nsun2-depletion in *Apc^fl/fl^* mouse small intestinal organoids by two best efficient *Nsun2*-targeting plasmids (*shNsun2-1 and shNsun2-2*). For *Nsun2* knockdown in *Apc^fl/fl^* organoids; transduction started on Day1, and before puro-selection was imaged on Day2. Puromycin selection started on Day3, and after puro-selection was imaged on Day6. Clonogenicity was performed on Day9 with 10,000 single cells, and Resazurin (24h) was measured on Day12. **F** The percentage of raw clonogenicity levels in *shNsun2-1* and *shNsun2-2* groups compared to *shCtrl* in *Apc^fl/fl^* mouse small intestinal organoids (data are presented as mean ± SD; ****p<0.0001, , Ordinary One-way ANOVA test compared to *shCtrl,* n = 5 experimental replicates). **G** The percentage of normalised organoid area (µm^2^) in *shNsun2-1* and *shNsun2-2* compared to *shCtrl* in *Apc^fl/fl^* mouse small intestinal organoids (data are presented as mean ± SD; **p:0.0065 (*shCtrl* vs *shNsun2-1*), **p:0.0034 (*shCtrl* vs *shNsun2-2*), Ordinary One-way ANOVA test compared to *shCtrl,* n = 4 experimental replicates). **H** The percentage of relative viability by Resazurin assay in *shNsun2-1* and *shNsun2-2* compared to *shCtrl* in *Apc^fl/fl^* mouse small intestinal organoids (data are presented as mean ± SD; **p:0.0003 (*shCtrl* vs *shNsun2-1*), **p:0.0002 (*shCtrl* vs *shNsun2-2*), Ordinary One-way ANOVA test compared to *shCtrl,* n = 4 experimental replicates). **I** mRNA expression levels of stem cell markers *(Lgr5, Ascl2, Igfbp4, Lrig1, Smoc2)* verifies the knockdown and shows the reduction in ISC signature, respectively. All mRNA expressions were normalised to *β-actin*, a housekeeping control. All data are presented using Ordinary one-way ANOVA compared to control statistical test. n≥3 experimental replicates

### *Nsun2* is required for *Apc*-loss mediated hyper-proliferation and stem cell expansion *in vivo*

To determine the role of NSUN2 in normal intestinal homeostasis and *Apc*-loss induced hyper-proliferation, we generated *Vil-CreERT2 Nsun2^fl/fl^*(*Nsun2*), and *Vil-CreERT2-Apc^fl/fl^ Nsun2^fl/fl^*(*Apc Nsun2*) mice by crossing to *Vil-CreERT2-Wt* (*Wt*) and *-Apc^fl/fl^* (*Apc*) lines^53^ (Fig 4A). Mice were induced with tamoxifen, intestinal crypts isolated 4 days post induction, and *Nsun2* knockout validated at mRNA and protein levels (Fig 4B-C). Immunohistochemistry staining also demonstrated effective deletion of NSUN2 in *Nsun2* and *Apc Nsun2* intestines (Fig 4D, bottom). Histological analysis of intestines revealed a reduction in *Apc*-loss driven hyperproliferation following *Nsun2* deletion with a significantly reduced number of BrdU^+ve^ proliferative cells in *Apc Nsun2* intestines compared to *Apc* (Fig 4D middle and Fig 4E). Additionally, expression of ISC markers, such as *Lgr5, Ascl2, Smoc2 and Lrig1* was substantially diminished in *Apc Nsun2* intestines (Fig 4F, Supplementary Fig S4A) consistent with our findings in in *Nsun2*-depleted *Apc^fl/fl^* organoids (Fig 3). To further validate the stem cell loss phenotype, we cultured single intestinal cells from *Apc* and *Apc Nsun2* intestines *ex vivo.* Cells isolated from *Apc Nsun2* intestines had reduced clonogenic capacity indicative of reduced stem cell function (Fig 4G-H and Supplementary Fig S4B).

**Figure 4.**
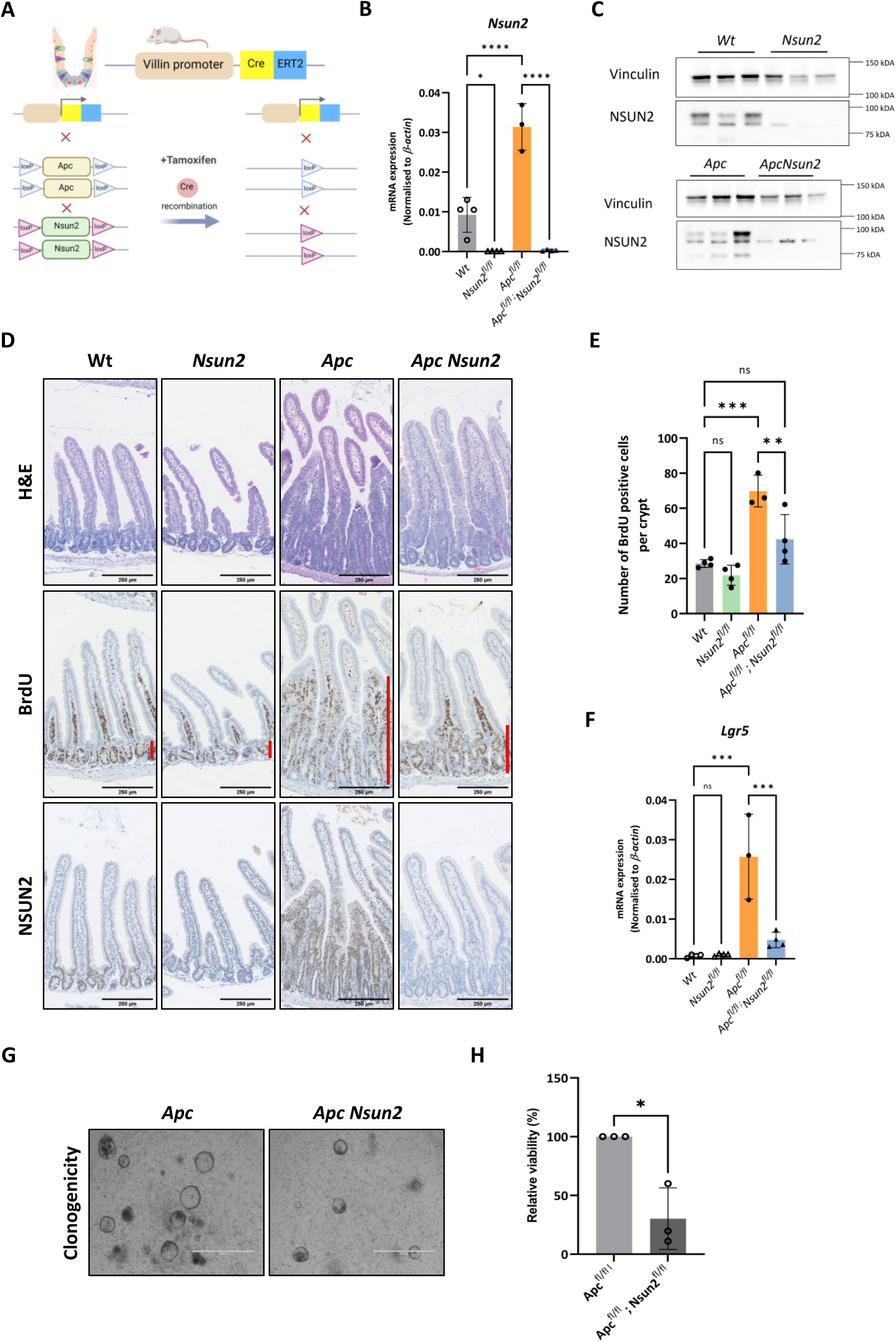
NSUN2 deletion reduces intestinal hyperproliferation and stem cell expansion following Apc deletion in mouse small intestine. **A** Schematic demonstration of *Apc* and *Nsun2* knockout mouse model generation in the intestinal tissue-specific Vil-CreERT2 line. The figure was created by Biorender.com. **B** Nsun2 mRNA expression and knockout validation of *Nsun2* deletion in Vil-CreERT2 wild-type, *Apc^fl/fl^* and, *Apc^fl/fl^ Nsun2^fl/fl^* mouse small intestines by qPCR. All mRNA expressions were normalised to *β-actin*, a housekeeping control (data are presented as mean ± SD; ***p:0.0002, ****p<0.0001, *p:0.03; One-way ANOVA test comparing the groups each other, n = minimum 3 biologically independent mice per group) **C** Validation of NSUN2 and a house-keeping control VINCULIN protein expressions in Vil-CreERT2 mouse *Apc^fl/fl^* and *Apc^fl/fl^ Nsun2^fl/fl^* mouse small intestinal tissues by western blot (n = 3vs3 biologically independent mice). **D** Representative images of Haematoxylin and Eosin (H&E), NSUN2, proliferating cells (BrdU), and β-CATENIN staining in wild-type, *Apc^fl/fl^* , *Apc^fl/fl^ Nsun2^fl/fl^* Vil-CreERT2 mouse small intestines. Orange lines indicate extension of crypt regions of small intestine. **E** Quantification of number of BrdU positive cells per crypt using QuPath program. (data are presented as mean ± SD; ***p:0.0004 (*Vil-Wt* vs *Vil-*, *Apc^fl/fl^*), **p:0.009 (*Vil-*, *Apc^fl/fl^ vs Vil-Apc^fl/fl^Nsun2^fl/fl^* ; One-way ANOVA test comparing the groups each other, n = minimum 3 biologically independent mice per group). **F** mRNA expression levels of stem cell marker *Lgr5* showing the reduction in ISC signature, respectively. All mRNA expressions were normalised to *β-actin*, a housekeeping control (data are presented as mean ± SD; ***p:0.0001, ***p:0.0001, and ***p:0.0005; One-way ANOVA test comparing the groups each other, n = minimum 3 biologically independent mice per group). **G-H** Representative images of clonogenicity assay (left) performed in direct crypt culture from *Apc^fl/fl^*, *Apc^fl/fl^ Nsun2^fl/fl^* Vil-CreERT2 mouse small intestines. 50,000 single cells seeded on Day 1 and relative cell viability (%) (right) was measured by 24h Resazurin incubation on Day 6 (data are presented as mean ± SD; *p:0.0438; Unpaired t-test with Welch’s correction, n = minimum 3 biologically independent mice per group).

Next, we investigated the effects of *Nsun2*-loss on normal intestine both at phenotypic and transcriptomic levels. Despite robust *Nsun2* deletion in *Nsun2* mice (Fig 4B-C), ISC gene expression was unchanged compared to *Wt* control mice (Fig 4F, Supplementary Fig S4A). Moreover, loss of *Nsun2* did not affect normal intestinal histology or proliferation (Fig 4D-4E). Together, these findings show that while *Nsun2* is dispensable for normal intestinal homeostasis, it is critical for sustaining *Apc-*loss driven stem cell expansion and hyperproliferation.

### Loss of *Nsun2* in *Apc*-deficient intestines reduces ISC signature gene expression and impairs stem cell transformation

To evaluate *Nsun2*-dependent transcriptomic changes during *Apc*-driven hyper-proliferation, we analysed *Apc* and *Apc Nsun2* intestinal crypts using RNA-seq. In total, we identified 1639 differentially expressed genes. Of these, 1055 genes were downregulated (log_2_FC≤-1, p_adj._≤0.05) whereas 584 genes were upregulated (log_2_FC≥1, p_adj._ ≤0.05) (Fig 5A and Table S1). Gene ontology analysis of downregulated genes found dysregulation of translation regulation, RNA processing and cell cycle regulation in *Nsun2*-loss *Vil-Apc* intestinal crypts (Fig 5B, left). Cell apoptosis and immune activation were among the enriched upregulated processes in *Apc Nsun2-*deficient mice (Fig 5B, right). We furthered the analysis of this dataset using gene set enrichment analysis (GSEA). This revealed a notable enrichment in gene-sets linked with intestinal stem cell signatures and Wnt signalling targets being suppressed in *Apc Nsun2* crypts, further supportive of ablated cancer stem cell function (Fig 5C). Additional analysis demonstrated and verified this negative regulation of intestinal stem cell markers and Wnt target genes following *Nsun2* deletion (Fig 5D). To determine the transcriptional effects of *Nsun2* deletion on normal intestine we also carried out RNAseq on *Wt* and *Nsun2* intestinal crypts. Consistent with the lack of phenotypic impact of *Nsun2* deletion on normal intestine we found far fewer transcriptional changes than in *Apc-*deficient tissue with only 4 genes downregulated and 7 genes upregulated (log_2_FC≥1, p_adj._ ≤0.05) (Table S2). Furthermore, we found no changes in expression of ISC markers or Wnt signalling targets, consistent with *Nsun2* being dispensible for normal intestinal homeostasis (Supplementary Figure S5A).

**Figure 5.**
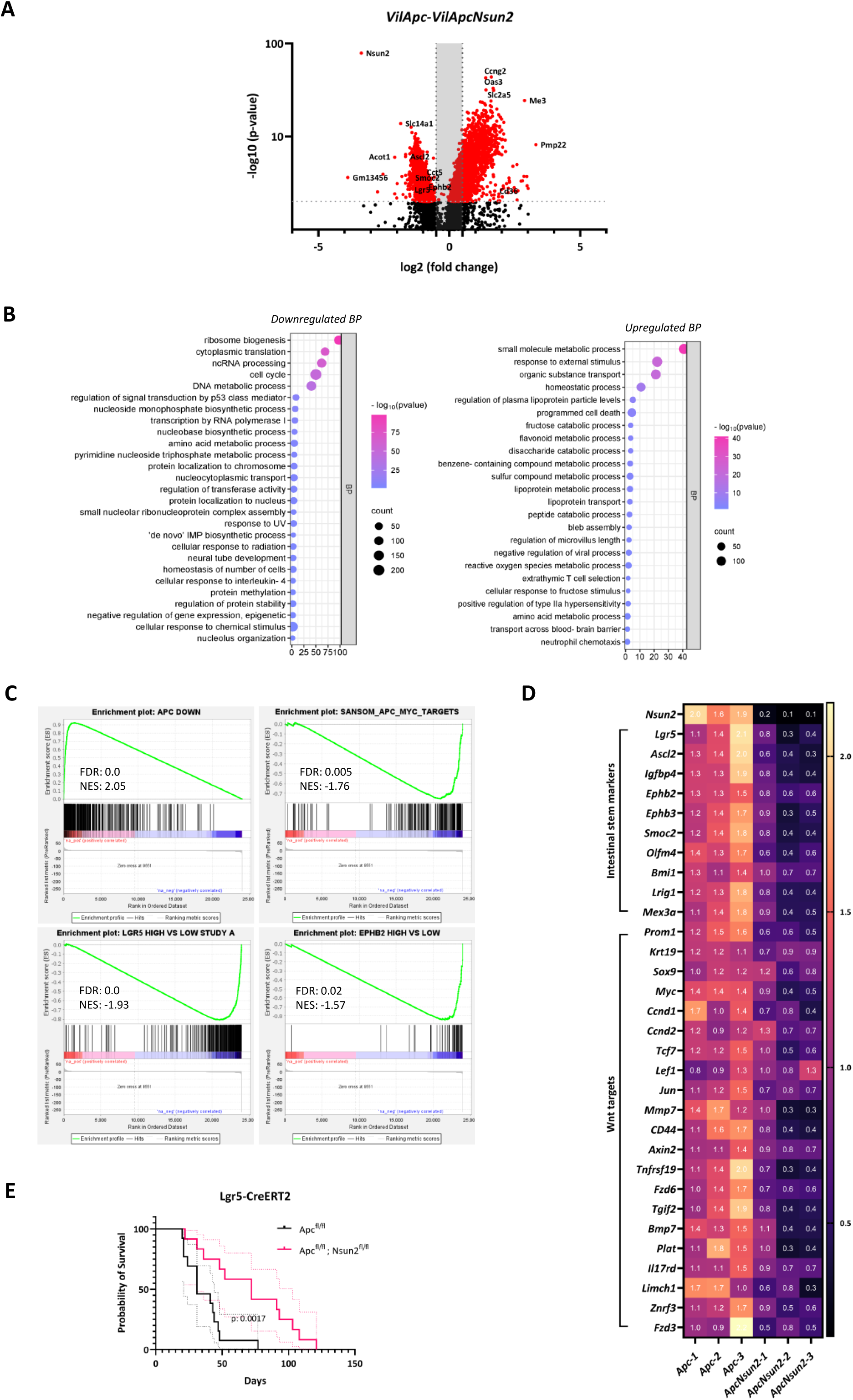
*Nsun2-loss* reduces stem cell signature along with Wnt-associated transcriptomic targets and improves mice survival. **A** The volcano plot shows the number of differentially expressed genes in *Vil-Apc* and *Vil-ApcNsun2* comparison. Differentially expressed genes (DEGs) were defined using log_2_FC±1 and p_adj._ ≤0.05. **B** Gene ontology analysis for biological process enrichment using downregulated (left) and upregulated (right) differentially expressed genes from RNA-seq comparison in *Vil-Apc* vs *Vil-ApcNsun2* mouse small intestinal crypts. GO plots are generated using SRplot^70^. **C** Gene set enrichment analysis (GSEA)-enrichment plots of representative gene sets comparing *Vil-Apc* vs *Vil-ApcNsun2* mouse small intestinal crypts. Negative enrichment score (NES) and FDR q values were shown in each panel. **D** Heat-map of differentially expressed intestinal stem cell markers and Wnt targets comparing small intestinal crypts from *Vil-Apc* vs *Vil-ApcNsun2* mice (n = 3 vs 3 biologically independent mice). **E** Probability of survival of *Lgr5-Apc* and *Lgr5-ApcNsun2* intestinal cohorts in *Lgr5-IRES-eGFP-CreERT2* mouse model. (data is presented as mean ± SD; **p:0.0017, as determined by Log-rank Mantel-Cox test, n=13 vs 12).

Due to the robust reduction in the ISC signature observed in *Nsun2*-deficient *in vitro* and *in vivo Apc^fl/fl^* tumour models, we next aimed to test whether NSUN2 is required for malignant stem cell transformation. To assess this, we used the *Lgr5-IRES-eGFP-CreERT2* model to induce intestinal adenomatous tumour formation from intestinal stem cells^2,4,54^ (Supplementary Figure S5B). Cohorts of *Lgr5-IRES-eGFP-CreERT2 Apc^fl/fl^* (*Lgr5 Apc*) *Lgr5-IRES-eGFP-CreERT2 Apc^fl/fl^ Nsun2^fl/fl^* (*Lgr5 Apc Nsun2*) were generated and tamoxifen administered to induce gene deletion. We found *Apc* loss driven ISC transformation and adenoma formation was significantly impaired by *Nsun2* deletion (Fig 5E). Together, these findings define a key role for NSUN2 in regulating ISC transformation and intestinal tumour initiation.

### NSUN2 mediates methylation of mRNAs involved in RNA processing, stem cell fate and oncogenic MAPK/ERK signalling

To identify the downstream mRNA targets of NSUN2-mediated m^5^C methylation in intestinal homeostasis and tumourigenesis, we performed whole transcriptome mRNA bisulphite sequencing (mRNA Bis-seq). We analysed both *in vivo* mouse intestinal crypts (*Vil Wt, Vil Nsun2, Vil Apc* and *Vil Apc Nsun2*) and *in vitro Apc^fl/fl^* organoids (*shCtrl* and *shNsun2*) (Fig 6A, Table S3-8). Global mRNA m^5^C analysis revealed significant changes in m^5^C methylation in our experimental groups (Supplementary Fig S6A). Focusing on the *Apc*-deficient models, we identified 13,374 and 11,134 differentially methylated sites (DMSs) in *Apc* vs *Apc Nsun2* intestinal crypts and *Apc^shCtrl^ vs Apc^shNsun2^* comparisons, respectively (Supplementary Fig S6A). Interestingly, methylation changes were bi-directional, with both increased and decreased methylation observed across DMSs (Supplementary Fig S6A). To identify common m^5^C sites in the two experimentally independent groups, we intersected the differentially methylated sites from *in vivo* intestinal crypts (*Apc* vs *Apc Nsun2*) and 3D organoids (*shCtrl* vs *shNsun2*). We detected 525 common m^5^C sites (hereafter termed *in vivo-vs-in vitro*) (Fig 6B) that either consistently lost methylation (295 sites) or consistently gained methylation (230 sites) (Table S9). Mapping these methylation changes to genomic region found that sites losing methylation were enriched in gene coding regions (223/525, 43%) whilst those gaining methylation mapped predominantly to mitochondrial gene loci (Figure 6C and Table S9).

**Figure 6.**
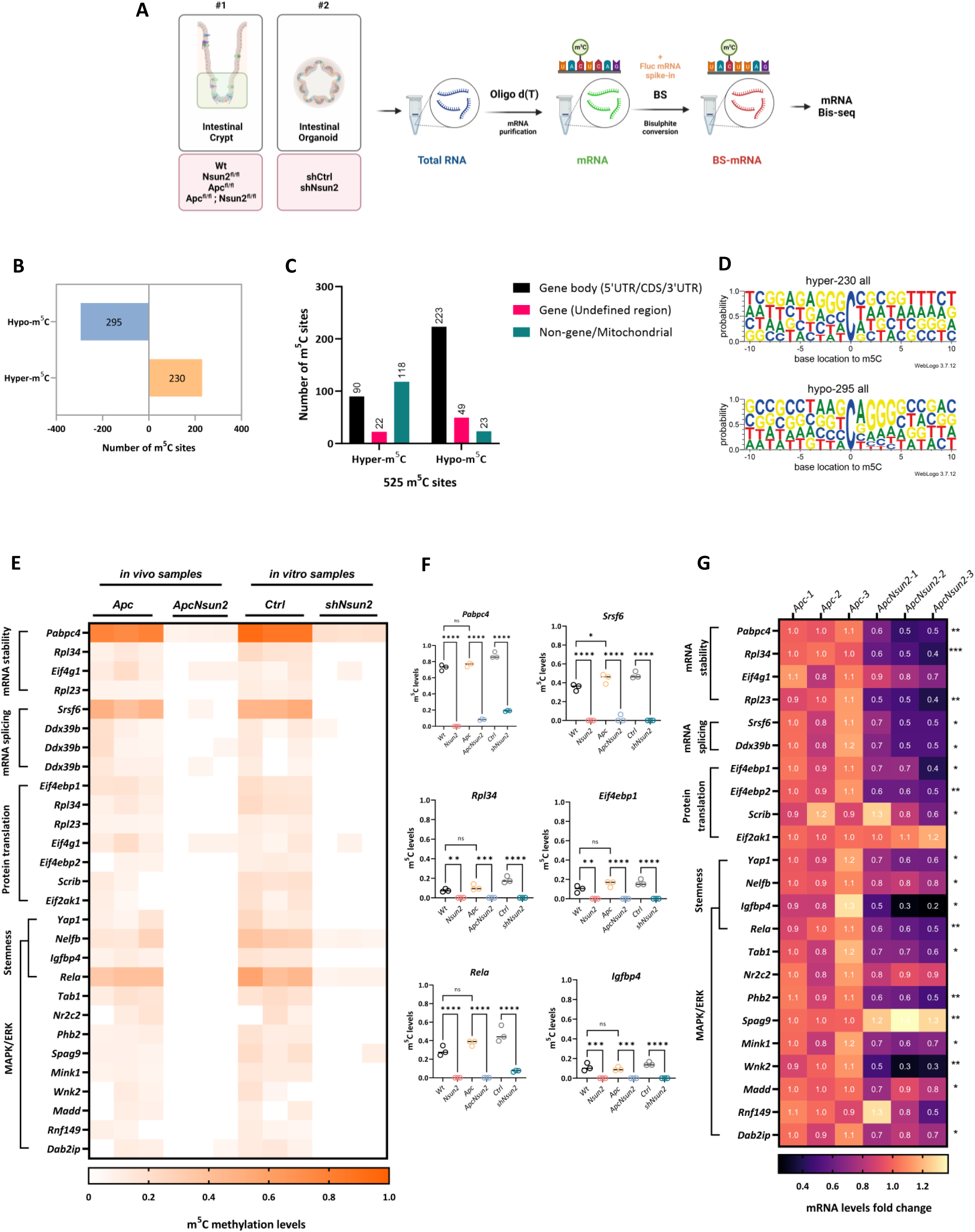
*Nsun2*-loss leads to bi-directional transcriptomic m^5^C signature in *Apc-deficient* intestines. **A** Schematic demonstration of experimental models used in whole transcriptome bisulphite sequencing and workflow. The figure was created by Biorender.com. **B** The bar graph shows the number of m^5^C sites that are common methylation gain and loss between *Vil-Apc* vs *Vil-ApcNsun2* and *shCtrl vs shNsun2* comparison. Differentially methylated sites (DMSs) were defined using Δmethylation ratio: ±0.05. **C** Bar graph showing the number of hyper and hypo methylated sites identified in different genomic regions. **D** The probability pattern represents sequence context of 10 bases up and downstream of hypo-methylated and hyper-methylated of common 525 NSUN2-dependent m^5^C sites as a result of overlapping vivo and vitro m^5^C sites **E** Heat-map of differentially methylated sites in mRNA stability, mRNA splicing, Protein translation, Stemness, and MAPK/ERK cascade in *Vil-Wt* vs *Vil-Nsun2, Vil-Apc* vs *Vil-ApcNsun2* intestinal crypts, and *shCtrl vs shNsun2* organoids (n = 3 vs 3 biologically independent mice or experimental replicates). **F** mRNA m^5^C methylation levels in some of the selected mRNAs involved in mRNA stability (*Pabpc4, Rpl34*), mRNA splicing (*Srsf6*), Protein translation (*Eif4ebp1*), Stemness and MAPK/ERK cascade (*Rela* and *Igfbp4*) in *Vil-Wt* vs *Vil-Nsun2, Vil-Apc* vs *Vil-ApcNsun2* intestinal crypts and *shCtrl vs shNsun2* by mRNA-bis-seq (data are presented as mean ± SD; ***p<0.001, **p<0.01, *p<0.05; One-way ANOVA test comparing the groups each other, n = 3 vs 3 biologically independent mice or experimental replicates). **G** Heat-map of mRNA levels in mRNA stability, mRNA splicing, Protein translation, Stemness, and MAPK/ERK cascade in *Vil-Apc* vs *Vil-ApcNsun2* intestinal crypts by RNA-seq (***p<0.001, **p<0.01, *p<0.05; Unpaired t-test, n = 3 biologically independent mice per group).

WebLogo3^55^ motif analysis revealed enrichment of a G**C**[A/C]GGG motif in common hypo-methylated sites (Fig 6D). This motif has previously been linked to NSUN2-mediated methylation, suggesting a conserved recognition sequence for NSUN2 activity^31,32,56^. Further analysis of mRNA regions showed that methylation sites altered following *Nsun2* depletion were enriched in the coding sequence (CDS; 119/525, 23%) and 3’ untranslated region (3’UTR; 135/525, 26%) although some were also found within the 5’ untranslated region (5’ UTR; 35/525, 7%) (Supplementary Fig 6B). Across different mRNA regions, the same G**C**[A/C]GGG enriched motif was observed in sites losing methylation, suggesting this is the consensus sequence for NSUN2 activity, regardless of DMS location on the mRNA molecule (Supplementary Fig 6C). Taken together, these data suggest NSUN2 activity is primarily directed towards specific, G rich regions in gene encoding mRNAs, with other methyltransferases likely responsible for methylation of mitochondrial transcripts.

To further understand the functional impact of NSUN2 dependent m^5^C mRNA methylation we investigated the function of hypomethylated mRNAs. This revealed targets of NSUN2 methylation involved in multiple cellular processes including RNA processing, stem cell function and oncogenic MAPK/ERK signalling (Fig 6E). Regarding RNA processing, *Nsun2* depletion led to hypomethylation of genes involved in mRNA stability (*Pabpc4*), mRNA splicing (*Srsf6, Ddx39b*) and protein translation (*Rpl23, Rpl34, Eif4g1, Eif4ebp1* and *Eif4ebp2*). We also observed hypomethylation of genes involved stem cell regulation including *Nelfb* which controls embryonic stem cell fate^57^ and *Rela* and *Yap1* which are previously described mediators of Wnt-driven ISC expansion and tumour initiation^4,58–60^. We also observed hypomethylation of numerous genes involved in oncogenic MAPK/ERK signalling including *Igfbp4, Rela, Tab1, Spag9, Mink1, Wnk2,* and *Dab2ip.* In addition to *Apc Nsun2* deficient models, loss of methylation in several of these transcripts was also observed in normal intestine following *Nsun2* deletion, further supporting their regulation by NSUN2 (Fig 6F). Analysis of mRNA expression levels demonstrated that these hypo-methylated transcripts were also downregulated in *Apc Nsun2* deficient intestinal crypts compared to *Apc*-deficient controls (Fig 6F, Table S10). Together, this indicates NSUN2 plays a key role in regulating the methylation status of mRNAs involved in RNA processing, stem cell activity and oncogenic signalling cascades, such as MAPK/ERK. Loss of m^5^C methylation in these pathways leads to reduced mRNA expression, likely contributing to the impaired stem cell function and tumour initiation observed in *Apc-*deficient tissue following *Nsun2* depletion.

### NSUN2 modulation of MAPK/ERK signalling mediates its oncogenic functions

As NSUN2-mediated m^5^C methylation affects multiple genes involved in the MAPK/ERK signalling pathway, we next assessed whether MAPK/ERK activity is altered in cells lacking NSUN2. First, we assessed the levels of phosphorylated-ERK1/2 (p-44,42), the active form of ERK, in *Apc^shNsun2^*organoids. We found significantly reduced p-ERK1/2 levels compared to *Apc^shCtrl^*(Fig 7A-B). Similarly, *in vivo* analysis of *Apc Nsun2* mouse intestinal crypts showed a significant reduction in p-ERK1/2 levels compared to *Apc* controls (Fig 7C-D).

**Figure 7.**
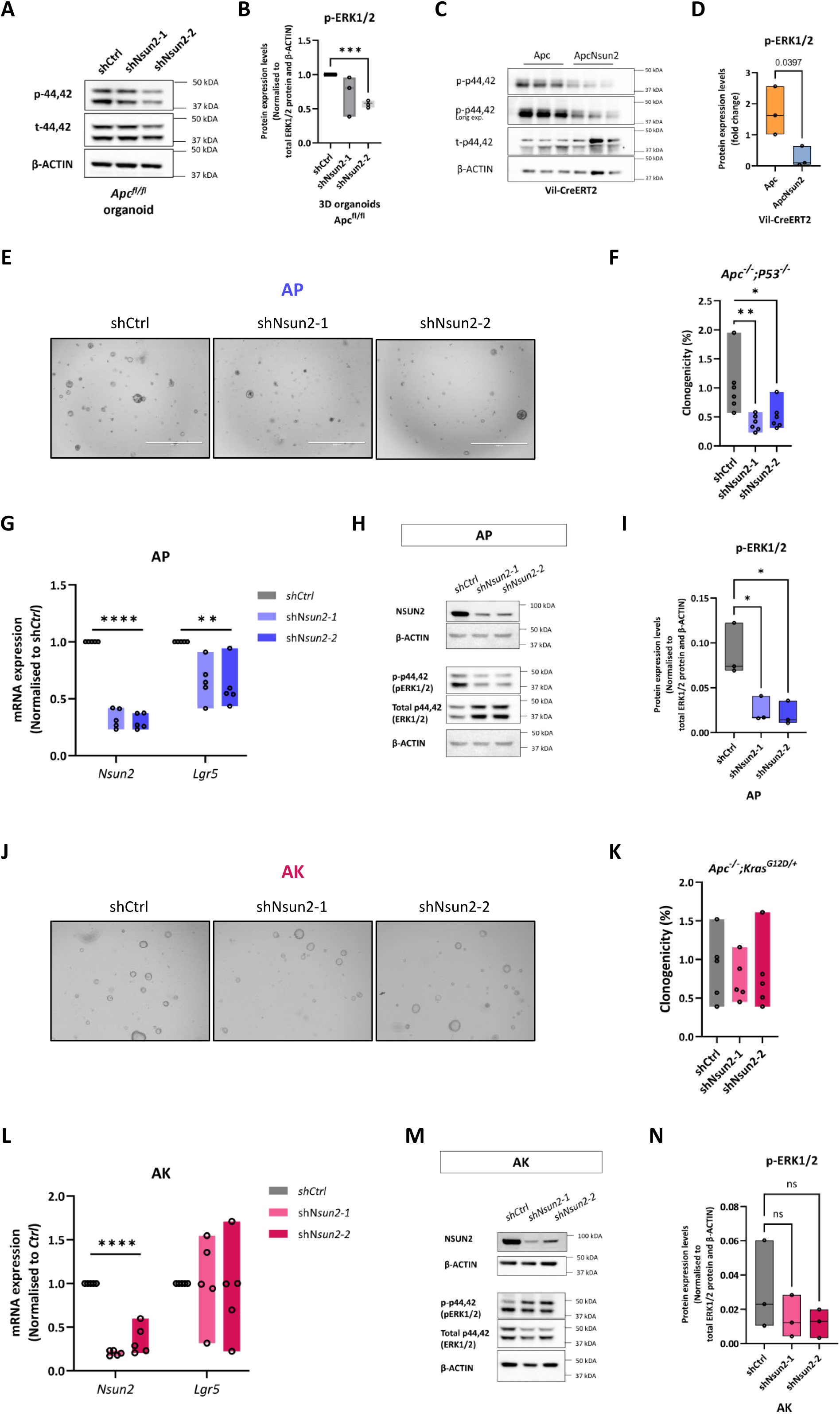
NSUN2 mediates MAPK activation. **A** Representative image of validation of Nsun2 mediating MAPK/ERK signalling through phospho-ERK1/2 (p-44,42) in *Apc^fl/fl^-shCtrl* and *Apc^fl/fl^-shNsun2* mouse small intestinal organoids by western blot. **B** Quantification of normalised phospho-ERK1/2 (p-44,42) protein expression levels in *Apc^fl/fl^-shCtrl* and *Apc^fl/fl^-shNsun2* mouse small intestinal organoids by western blot, using β-ACTIN loading control (data are presented as mean ± SD; ***p:0.0001, two-tailed t-test, n = 3 experimental replicates). **C** Representative image of validation of Nsun2 mediating MAPK/ERK signalling through phospho-ERK1/2 (p-44,42) in *Vil-Apc* vs *Vil-ApcNsun2* intestinal crypts by western blot. **D** Quantification of normalised phospho-ERK1/2 (p-44,42) protein expression levels in *Vil-Apc* vs *Vil-ApcNsun2* intestinal crypts by western blot, using β-ACTIN loading control (data are presented as mean ± SD; *p:0.0397, two-tailed t-test, n = 3vs3 biologically independent mice). **E** Representative image of Nsun2-depletion in *Apc^fl/fl^ ; P53^fl/fl^ (AP)* mouse small intestinal organoids by two best efficient *Nsun2*-targeting plasmids (*shNsun2-1 and shNsun2-2*). For *Nsun2* knockdown in *AP* organoids; transduction started on Day1, and before puro-selection was imaged on Day2. Puromycin selection started on Day3, and after puro-selection was imaged on Day6. Clonogenicity was performed on Day9 with 2,000 single cells, and Resazurin (24h) was measured on Day12. **F** The percentage of raw clonogenicity levels in *shNsun2-1* and *shNsun2-2* groups compared to *shCtrl* in *Apc^fl/fl^ ; P53^fl/fl^ (AP)* mouse small intestinal organoids (data are presented as mean ± SD; **p:0.0065 and *p:0.023, Ordinary One-way ANOVA test compared to *shCtrl*, n = 6 experimental replicates) **G** Validation of *Nsun2* knockdown and mRNA expression level stem cell marker gene *Lgr5* in *Apc^fl/fl^* mouse small intestinal organoids by qPCR. All mRNA expressions were normalised to *β-actin*, a housekeeping control (data are presented as mean ± SD; ****p<0.0001, **p:0.0073 and **p:0.0032, Ordinary One-way ANOVA test compared to *shCtrl*, n = 5 experimental replicates). **H** Representative image of validation of NSUN2 depletion and NSUN2 mediating MAPK/ERK signalling through phospho-ERK1/2 (p-44,42) in *Apc^fl/fl^ ; P53^fl/fl^ (AP)* mouse small intestinal organoids following Nsun2-knockdown intestinal crypts by western blot. β-ACTIN used as loading control. **I** Quantification of normalised phospho-ERK1/2 (p-44,42) protein expression levels in *Apc^fl/fl^ ; P53^fl/fl^ (AP)-shCtrl* and *Apc^fl/fl^ ; P53^fl/fl^ (AP)-shNsun2* mouse small intestinal organoids by western blot, using β-ACTIN loading control (data are presented as mean ± SD; *p:0.0152 and *p:0.0113, Ordinary One-way ANOVA test compared to *shCtrl*, n = 3 experimental replicates). **J** Representative image of Nsun2-depletion in *Apc^fl/fl^ ; Kras^G12D/+^ (AK)* mouse small intestinal organoids by two best efficient *Nsun2*-targeting plasmids (*shNsun2-1 and shNsun2-2*). For *Nsun2* knockdown in *AP* organoids; transduction started on Day1, and before puro-selection was imaged on Day2. Puromycin selection started on Day3, and after puro-selection was imaged on Day6. Clonogenicity was performed on Day9 with 2,000 single cells, and Resazurin (24h) was measured on Day12. **K** The percentage of raw clonogenicity levels in *shNsun2-1* and *shNsun2-2* groups compared to *shCtrl* in *Apc^fl/fl^ ; Kras^G12D/+^ (AK)* mouse small intestinal organoids (data are presented as mean ± SD; not significant values are not indicated, Ordinary One-way ANOVA test compared to *shCtrl*, n = 5 experimental replicates). **L** Validation of *Nsun2* knockdown and mRNA expression level stem cell marker gene *Lgr5* in *Apc^fl/fl^ ; Kras^G12D/+^ (AK)* mouse small intestinal organoids by qPCR. All mRNA expressions were normalised to *β-actin*, a housekeeping control (data are presented as mean ± SD; ****p<0.0001, not significant values are not indicated, Ordinary One-way ANOVA test compared to *shCtrl*, n = 5 experimental replicates). **M** Representative image of validation of NSUN2 depletion and NSUN2 mediating MAPK/ERK signalling through phospho-ERK1/2 (p-44,42) in *Apc^fl/fl^ ; Kras^G12D/+^ (AK)* mouse small intestinal organoids following Nsun2-knockdown intestinal crypts by western blot. β-ACTIN used as loading control. **N** Quantification of normalised phospho-ERK1/2 (p-44,42) protein expression levels in *Apc^fl/fl^ ; Kras^G12D/+^ (AK)-shCtrl* and *Apc^fl/fl^ ; Kras^G12D/+^ (AK)-shNsun2* mouse small intestinal organoids by western blot, using β-ACTIN loading control (data are presented as mean ± SD; Ordinary One-way ANOVA test compared to *shCtrl*, n = experimental replicates).

To confirm these findings, we next used an *Apc^fl/fl^;P53^fl/fl^*(*AP*) organoid model, which allows investigation of the combined effects of *Apc* loss and *P53* inactivation (which commonly occurs during CRC progression). Similar to *Apc^fl/fl^* organoids, depletion of *Nsun2* using shRNA in *AP* organoids (Supplementary Fig 7A) led to diminished clonogenicity, number and area of organoid formation in addition to viability (Fig 7E-F and Supplementary Fig 7B). In accordance with decreased colony formation ability, expression of the stem cell marker gene, *Lgr5*, is also reduced in *AP^shNsun2^*organoids (Fig7G). Consistent with the *Apc* models, *Nsun2* depletion in AP organoid resulted in significantly reduced ERK1/2 phosphorylation (Fig 7H-I), confirming that NSUN2 regulates ERK activation in *Apc*-deficient CRC models.

To functionally test the role of MAPK/ERK signalling, we tested whether constitutive activation of the pathway via *Kras^G12D^* mutation could bypass the requirement for NSUN2 in tumourigenesis. Using the *Apc^fl/fl^ Kras^G12D/+^* (*AK*) organoid model, we found that the effects of *Nsun2* depletion on colony formation ability, stem cell marker expression, cell viability and p-ERK1/2 activation was reversed by *Kras^G12D/+^* mutation (Fig 7J-M and Supplementary Fig 7C-D). This suggests that the *AK* organoids, through the *Kras^G12D/+^* mutation, rewire the MAPK/ERK pathway, resulting in constitutive activation of MAPK/ERK signalling (Fig 7M-N). Together, these findings demonstrate that NSUN2 promotes ISC expansion and CRC initiation via regulation of MAPK/ERK signalling.

## Discussion

In this study, we demonstrate that NSUN2 is a critical mediator of Wnt-driven intestinal stem cell transformation. Depletion of *Nsun2* impairs growth and stem cell function in *Apc*-deficient *in vitro* organoids and *in vivo* intestines while leaving healthy tissue unaffected. Furthermore, *Nsun2*-depletion suppresses tumourigenesis in a stem cell driven model CRC. Transcriptome-wide analysis reveals NSUN2-dependent mRNA methylation changes, particularly affecting stem cell regulators and components of the MAPK/ERK pathway. Notably, MAPK pathway activation rescues stem cell dysfunction caused by *Nsun2* loss, highlighting NSUN2’s role in CRC initiation via MAPK/ERK regulation.

Previous studies have linked NSUN2 to MYC-driven proliferation and tumourigenesis in various cancers^27,37,45,56,61^. Consistent with work reported by Frye and Watt, which showed overexpression of NSUN2 in a limited number of CRC patients^37^, we observed elevated NSUN2 in *Apc*-driven hyper-proliferative and malignant intestinal tissues. Increased NSUN2 expression has been reported across multiple malignancies, such as gastric^45^, oesophageal^47^, hepatocellular^46^, bladder cancer^49^ where it correlates with poor prognosis^26,45,47,56^. Functional parallels also exist between our findings and previous studies. For example, Blanco et al. revealed that NSUN2-loss disrupts epidermal stem cell function and cell cycle progression, along with dysregulating site-specific tRNA methylation^36,62^. Similarly, we observe reduced expression of intestinal CSC markers, including *Lgr5*^3^, *Ephb2*^63^, *Lrig1, Smoc2* and *Igfbp4*. Yet, despite a clear stem cell dysfunction following *Apc* deletion we find no requirement for NSUN2 function in healthy intestinal stem cells.

Using bisulphite mRNA sequencing, we identified NSUN2-dependent mRNA methylation sites in both organoids and tissues. Comparative analysis of these datasets identified 525 common regulated sites in these models. Due to the phenotypic similarity of *Nsun2* depletion *in vitro* and *in vivo* these likely represent robust, functionally relevant m^5^C methylation changes. Interestingly, we observed both losses and gains of m^5^C methylation following *Nsun2* depletion. Whilst unexpected, this is similar to a recent study in the HCT15 colorectal cancer cell line showing *Nsun2*-dependent losses and gains of m^5^C instead of a global m^5^C reduction^64^. One possibility is the presence of other, compensating m^5^C methyltransferases or a more complex interplay of m^5^C regulation than previously anticipated. Analysis of *Nsun2* dependent methylation sites led to identification of an enrichment of G**C**[A/C]GGG dominant motif, suggesting a role for CG content in guiding NSUN2 activity. We observe this motif in all mRNA regions targeted by NSUN2, in line with previously reported slightly different type I motif of m^5^C, which contains a downstream G-rich triplet that have been reported in 5’UTR, CDS, and 3’UTR^30^. Similar to this, another study reported the N1-methyladenosine (m^1^A) modification occurs in C/G-rich sequence context^65^. The functional significance of these consensus motifs remains unclear. However, it is known that CG content significantly influences RNA structure, which may, in turn, affect translation rates. This suggests that the CG content could play a crucial role in determining how efficiently proteins are synthesized from mRNA^65^. More broadly, these motifs could help predict site-specific RNA methylation or identify novel m^5^C regulators that specifically binds these regions.

We identified MAPK/ERK activation as a key pathway regulated by NSUN2. This is consistent with previous studies in esophageal cancer cell lines where *NSUN2* depletion suppresses activation of PI3K/AKT and MAPK/ERK signalling, a finding that parallels our observations in organoid and *in vivo* CRC models. *Apc^fl/fl^* organoids with *P53* deletion are also dependent on *Nsun2* expression for maintaining stem cell function and ERK signalling. However, *Kras^G12D/+^* mutation rescues the phenotypic effects of *Nsun2* loss suggesting *Kras^G12D/+^* sustains ERK signalling, downstream of NSUN2. Together, these findings demonstrate NSUN2 is a critical mediator of ISC expansion and tumour initiation via control of MAPK/ERK pathway activation. This defines a general mechanism by which RNA modifications mediate tumour initiation via maintenance of oncogenic signalling pathway activity.

## Methods

### Ethics statement

All mouse experiments were performed under the regulations by UK Home Office and all related ethical regulations were adhered to. The study protocols were approved by the University of Edinburgh AWERB. All procedures were carried out within the remits of both UK Home Office Project Licence (7008885) and Personal License (I97725097).

### Mouse experiments

All mice used in this project were housed at the Biomedical Research Facility (BRF) located at the Western General Hospital Campus of The University of Edinburgh. All mice were fed with a standard diet along with 12h light/dark cycle. When under experiment the mice were kept to a maximum of five per cage, but most commonly were housed in cages of two or three. Single housing of mice only used for the male mice separated from cage-mates due to being culled. However, single female mice were rehoused with other groups if possible. All mice were genotyped using Transnetyx (Cordoba, USA).

The alleles used in this study were as follows: Vil-CreERT2-Apc^fl^, C57BL/6N-Nsun2^tm1c(EUCOMM)Wtsi/WtsiOulu^ (Sperm) (Wellcome Trust Sanger Institute), and Lgr5-IRES-eGFP-CreERT2. To create a mixed line for co-deletion of these genes, the mice mentioned above were bred. After breeding, three different alleles (homozygote (fl/fl), heterozygote (fl/+), and wild-type (+/+)) are created for both *Apc* and *Nsun2* genes. Subsequent matings generate cohorts of appropriate combinations of *Apc* and *Nsun2* floxed alleles for experimental use. Then, Tamoxifen induction permits conditional tissue-specific *Apc* and/or *Nsun2* ablation following Villin or Lgr5 promoter. To simplify the explanation of the studies, the mice obtained from this model will be aforementioned as follows; Vil-CreERT2-wild-type (Vil-Wt), Vil-CreERT2-Nsun2^fl/fl^ (Vil-Nsun2), Vil-CreERT2-Apc^fl/fl^ (Vil-Apc), and Vil-CreERT2-Apc^fl/fl^;Nsun2^fl/fl^ (Vil-ApcNsun2), Lgr5-IRES-eGFP-CreERT2-Apc^fl/fl^ (Lgr5-Apc) Lgr5-IRES-eGFP-CreERT2-Apc^fl/fl^;Nsun2^fl/fl^ (Lgr5-ApcNsun2).

### Tamoxifen induction

For a short-term gene deletion; Cre recombination was induced in Villin-expressing epithelium by intraperitoneal injection (IP) of Tamoxifen (Sigma-Aldrich, T5648). Mice were given a 300 µL, 10 mg/mL dose of tamoxifen on day 0 and a 200 µL, 10 mg/mL dose of tamoxifen dose on day 1, and a 200 µL, 10 mg/mL dose of tamoxifen dose on day 2. Then, mice were humanely sacrificed by cervical dislocation (CD) in line with UK Home Office regulations and samples were collected on Day 5.

For the tumour cohort; Cre recombination was induced in Lgr5-expressing epithelium by oral gavage (OG) of tamoxifen. Mice were given a 200 µL, 10 mg/mL dose of tamoxifen dose for four consecutive days and aged. Mice were monitored, weighed, and assessed for symptoms at least twice a week. Then, mice were humanely sacrificed by CD in line with UK Home Office regulations depending on experiment and samples were collected. All mice given Tamoxifen irrespective of floxed alleles, rendering wild-type as a control group.

### In vivo labelling of cell proliferation

200uL of Bromodeoxyuridine (BrdU), an analogue of thymidine, (GE Healthcare, RPN201) was administered to mice by intraperitoneal injection for labelling of actively replicating cells 1 or 2-hour prior to humanely killing.

### Crypt isolation from mouse small intestine

Once mice were humanely killed by cervical dislocation, the first 10 cm of small intestinal (SI) tissue from mice was dissected and scraped carefully. Collected tissues were kept on ice until extraction started. SI was cut into small pieces and washed with cold PBS four times until the debris disappeared. Then, 100ul of 0.5M EDTA in 25ml cold PBS was added to the small intestine pellet and placed into a roller in a cold room for 30 min. This step loosens up the crypts from surrounding tissue. After incubation, 1:1 (v:v) Advanced DMEM/F12 (ADF) media was added on top of the EDTA-PBS solution. Then, the pellet was centrifuged at 300g for 5 min and washed once with cold PBS to remove excess EDTA. After this, 2nd, 3rd, and 4th fractions were collected by physical disruption using ADF media. These fractions were pooled and washed with ADF media once centrifuging at 500g for 5 min, then proceeded to either crypt culture or preservation in -80°C.

### 3D organoid models

Several colorectal cancer mouse organoid models (*Apc^fl/fl^, Apc^fl/fl^;Nsun2^fl/fl^*, *AK (Apc^fl/fl^;Kras^LSL-G12D/+^)*, and *AP (Apc^fl/fl^;Tp53^fl/fl^)* were generated via crypt isolation from corresponding mouse models or molecular methods by the members from Dr. Kevin Myant’s lab were used. Organoids were passaged depending on genotypes varying between 3-5 days and cultured in Cultrex PathClear Reduced Growth Factor BME (Bio-Techne, 3533–010–02). The plate was incubated for 10 min at 37 °C to allow BME solidification and 500 µl of growth medium was added. The growth medium contains Advanced DMEM/F-12 medium (ADF, Gibco) supplemented with, L-Glutamine and Penicillin/Streptomycin, Primocin, 1X B27 (Gibco, 17504044), 1X N2 (Gibco, 17502048), 50ng/ml EGF (Peprotech, 315–09–500), 100ng/ml Noggin (Peprotech, 250–38–500).

### shRNA knockdown

#### Plasmid constructs

Competent bacteria containing target gene set (Nsun2) plasmids (RMM3981- 201810604 and RMM3981-201837916) (AAATAAAGGATCATCTTCAGG and AAGTCTAGGTATGCTGGATGC) were ordered from Horizon discovery. Competent bacteria containing mouse Nsun2 shRNA plasmids as well as bacteria containing pLKO.1 plasmids (as a control) were grown in L-ampicillin plates overnight in 37°C incubator. Next day, single colonies were picked and cultured in liquid L-broth culture supplemented with L-ampicillin (50mg/ml) in 37°C shaker overnight. Subsequently, plasmids were isolated by QIAGEN Plasmid Midi (#200119) or Maxi Kit (#12662).

#### Lentiviral production

10 μg of a gene-specific lentiviral vector was mixed with 7.5 μg of the lentiviral packaging vector psPAX2 (Addgene) and 2.5 μg of the envelope protein-producing vector pCMV-VSV-G (Addgene). This mixture was then transfected into HEK 293T cells cultured in a 10 cm² dish using the polyethylenimine transfection reagent (Polysciences, 23966). After 48 hours, the virus was purified by filtering the supernatant media with a 0.45 μm filter, followed by concentration using the Lenti-X Concentrator (Takara Bio, 631232) and resuspension of the viral particles in PBS.

#### Lentiviral transduction of mouse colorectal cancer organoid models

Firstly, organoids were pre-treated with 1:10,000 Valproic Acid (VPA) on Day (-1). On Day 1, organoids were dissociated into single cells with 1ml pre-warmed StemPro Accutase for 2min at 37°C. The dissociation was stopped using 1ml of filtered 1% BSA in PBS. Resuspension was filtered and centrifuged at 300g for 3min at 4°C. The pellet resuspended in transduction media containing culture media and 8ug/ml Polybrene, 2μl Y27632 Rock Inhibitor (1:500), and Valproic acid (VPA) (1:10,000). The number of single cells was counted using automatic cell count (Countess II System). 100,000 cells containing 10μl virus per well was seeded into pre-coated plate with BME and incubated at 37°C for 24h. Media containing organoid-virus-polybrene sets on BME monolayer. On Day 2, media containing virus was removed, 100μl BME added per well. Once BME set, organoids were grown in culture media with 2μl Y27632 Rock Inhibitor (1:500). On Day 3, media was replaced with 2μg/ml Puromycin and incubated at 37°C for 72h. Puromycin selection kills non-transduced organoids. On Day 6, RNA and protein samples were isolated and reserved at -80°C or clonogenicity assay was performed. The organoids were incubated at 37°C, 0.5% CO^2^ throughout the experiments.

### Analysis of colony formation ability and metabolic activity by clonogenicity and Resazurin assay

The organoids were dissociated into single cells 6 days after the lentiviral transduction. 10,000 single cells (*Apc^fl/fl^*) and 2000 single cells (*AK* and *AP*) in 5μl BME were seeded to a 12-well plate. Sphere formation was observed following 2-4 days. Then, spheres were counted on Leica Light microscope and pictures on EVOS FL Cell Imaging System.

Once the organoid spheres were formed, the clonogenicity was counted and the media was replaced with a fresh media containing 10% Resazurin for 24h in 37°C incubator. Next day, relative cell viability was measured by multi-label counter Victor 21420 or TECAN Spark microplate reader in 96- well plate with three technical replicates.

### Total RNA and mRNA purification

RNA samples from mouse intestinal tissues, cell lines and organoid cultures were isolated by using RNeasy® Mini Kit following the manufacturer’s protocol. Subsequently, DNA degradation step was used by an additional step using RNase-Free DNase Set. RNA samples were analysed for quality and quantity using Thermo Scientific™ Nanodrop™ (Thermo Fisher).

To purify poly(A) mRNA, total RNA was subjected to oligo (dT)25 magnetic beads to capture poly(A) containing mRNA. The manufacturer’s protocol used with several adjustments. Firstly, 30–40 μg total RNA was incubated at 70°C for 5 min to disrupt secondary structures and placed on ice. Then, lysis/binding buffer suspensions were prepared according to manufacturer’s instruction. During the binding step, RNA samples added to Dynabeads/binding buffer were mixed for 20min at room temperature with gentle rotation at 5-10rpm for annealing of the mRNA to oligo (dT)25 magnetic beads. The mRNA-beads complex was washed twice before and eluted with 10 mM Tris-HCl at 70°C for 5 min. Finally, the enriched mRNA concentration was determined by Nanodrop, and the purity was analysed by Bioanalyzer Agilent RNA 6000 Nano/Pico assay (Agilent, Santa Clara, CA).

### cDNA synthesis and RT-qPCR

cDNA was generated by reverse transcription of the isolated RNA using qScript™ cDNA SuperMix with three RT-PCR cycles; 25°C for 5 min, 42°C for 30 min, 72°C for 15 min, 4°C for forever. qPCR performed according to the manufacturer’s protocol using fast SYBR Select Master Mix. Primer sequences are listed in Supplementary file. Amplifications were performed in two experimental replicates in 384-well plate using CFx Touch 384 (BioRad). β-Actin or Gapdh primers were used as a housekeeping control. Ct-values were normalized to β-Actin or Gapdh Ct -values. mRNA expression levels were calculated according to the ΔCt method and expressed as 2(- ΔCt).

### Protein isolation and quantification

Samples were washed with 1X cold PBS and centrifuged at 2000rpm for 5min. The pellet was resuspended with RIPA buffer (Sigma, #R0278) containing 10% (v/v) Proteinase and Phosphatase inhibitor cocktails and incubated on ice for 30-45min. Then, proteins were isolated after centrifuging at 14,000rpm at 4°C for 10min and isolated proteins were stored at -80°C. Isolated proteins were quantified using PierceTM BCA assay kit (Thermo Fisher).

### Western blot

Protein samples were prepared using 4X loading buffer (DTT) and denatured at 99°C for 5min. After taking immediately on ice, 15ug of proteins were loaded and run in 4-12% Bis-Tris or 3-8% Tris-Acetate gel at 150V for 1hour. Then proteins were transferred to nitrocellulose membrane by wet- western blotting technique. After transfer, membranes were stained in Ponceau S dye. Subsequently, the membranes were blocked, incubated with primary and corresponding HRP-linked secondary antibodies for each antibody as shown in Supplementary file. Finally, image visualisation was performed using chemiluminescent ECL substrates in Amersham ImageQuant 800. The quantification of the blots was performed using ImageJ68.

#### For stripping

the membranes were washed in 1X ReBlot Plus Strong Antibody Stripping Solution Reblot (Merck, #2504) for 10min and washed with PBST for 10min prior to re-blocking and re-incubation with new antibody. The membranes were stripped maximum two times.

### Dot blot

RNA samples were serially diluted (100ng - 300ng in total volume 4-7μl) and denaturised at 95°C for 5 min to disrupt secondary structures on Day 1. Then, RNA samples were immediately chilled on ice for 1 min to prevent re-formation of secondary structures and samples dropped onto Hybond-N+ membrane. The membrane was air-dry for 5min and RNA samples were crosslinked to the membrane by Stratalinker 2400 UV Crosslinker twice using the Autocrosslink mode (1,200 microjoules [x100]; 25-50 sec). Membrane was directly blocked with LICOR blocking buffer, incubated with primary and corresponding HRP-linked secondary antibodies for each antibody as shown in Supplementary file. Finally, image visualisation was performed using chemiluminescent ECL substrates in Amersham ImageQuant 800. The quantification of the blots was performed using ImageJ.

### Tissue sampling

Following from BrdU labelling and humane killing, small intestine and colon tissues were dissected from the mice. The gut was initially flushed with PBS to clean the lumen, opened longitudinally, pinned to flat surface, and fixed with paraformaldehyde for 20min at room temperature in the case of swiss-roll of the gut. Otherwise, three pieces of gut were cut and put together in a PFA containing tube, called parcel. Then, fixations continued overnight at 4°C. After 24hours, samples were transferred into 70% Ethanol. After fixation of the tissues, Tissue Tek® (Sakura) VIP processor was used to process the samples before being embedded in paraffin wax. After being embedded, 5μm sections of tissues were cut and dried overnight at 37°C incubator for further analysis, H&E or immunohistochemistry.

#### For Haematoxylin & Eosin (H&E) staining

Tissue sections on slides were dewaxed in three cycles of Xylene for 5 min in each, two cycles of 100% Ethanol, then one cycle of 75%, 50%, and 25% Ethanol for 2min in each, consecutively. Slides were then washed and incubated in haematoxylin for 90 seconds. After this, slides were washed with water for 2min, and left in lithium carbonate for 5 seconds to develop the signal, and washed quickly. Then, the slides were incubated in Eosin dye for 40sec to counterstain and washed again. Subsequently, slides were dehydrated by the opposite direction of dewaxing solutions and finally were mounted using DPX.

For standard IHC techniques and details, see Supplementary file. To scan the slides, Nanozoomer (Hammamatsu) and to image the slides NDP-image and QuPath softwares were used.

### RNA sequencing

For RNA sequencing, the quality of total RNA samples was initially assessed on the Agilent 2100 Electrophoresis Bioanalyser Instrument (Agilent Technologies Inc, #G2939AA) and RNA 6000 Nano chips (#5067-1511) at IGC Technical service. Then, the total RNA samples were sent to the Welcome Trust Clinical Research Facility (WTCRF) at the Western General Hospital, Edinburgh. All subsequent analysis for RNA sequencing were handled at WTCRF. The total RNAs were re-assessed using Qubit 2.0 Fluorimeter (Thermo Fisher Scientific Inc, #Q32866), Qubit RNA broad range assay kit (# Q10210), and the Qubit dsDNA HS assay kit (#Q32854) prior to library preparation. Then, libraries were prepared from 500ng of each total RNA sample using the NEBNEXT Ultra II Directional RNA Library Prep kit (NEB #7760) and the Poly(A) mRNA magnetic isolation module (NEB #E7490) according to the provided protocol. Subsequently, sequencing was performed on the NextSeq 2000 platform (Illumina Inc, #20038897) using NextSeq 2000 P2 Reagents (200 Cycles) (##20046812). Each library ended up with more than 24 million paired end reads. Raw FASTQ files were then uploaded for RNAseq analysis. To analyse differentially expressed genes, an interactive RaNA-seq tool was used. For functional analysis; g:Profiler, and gene set enrichment analysis; GSEA platforms were used.

### mRNA Bisulphite treatment

mRNA Bisulphite treatment was performed according to the manifacturer’s protocol in EZ RNA Methylation™ Kit. Only modification performed in mRNA conversion step by PCR are as follows; 70°C 10min, 64°C 45min, and 4°C.

### mRNA Bisulphite sequencing

The bisulphite-treated mRNA samples were assessed on Agilent 2100 Electrophoresis Bioanalyser Instrument using RNA 6000 Pico chips at IGC Technical service. Then, the samples were sent to WTCRF and were re-assessed by Qubit 2.0 Fluorimeter (Thermo Fisher Scientific Inc, #Q32855) and the Qubit RNA high sensitivity assay kit (# Q10210) prior to library preparation. After the quality control steps, the libraries were prepared from 10ng of each bisulphite-treated mRNA sample using the protocol 4 (library preparation for poly(A)-purified mRNA or rRNA-depleted RNA) NEBNEXT Ultra II Directional RNA Library Prep kit (NEB #7760). The mRNAs were fragmented. Subsequently, sequencing was performed on the NextSeq 2000 platform (Illumina Inc, #20038897) using NextSeq 2000 P3 Reagents (200 Cycles) (#20040560). Each library ended up with more than 46million paired end reads. Raw FASTQ files were then uploaded for the m^5^C-mRNA analysis.

### mRNA-Bis-seq analysis for 5-methylcytidine

Low-quality reads (Q<20) and adapter contaminations were removed by FASTQC and Trim Galore! tools on Galaxy (https://usegalaxy.org/). The argument in Trim Galore tool is to remove non-original sequence in the samples; read 1 ‘-j 1 -e 0.1 -q 20 -O 1-a AGATCGGAAGAGCACACGTCTGAACTCCAGTCAC’ and read 2 ‘-j 1 -e 0.1 -q 20 -O 1-a AGATCGGAAGAGCGTCGTGTAGGGAAAGAGTGTAGATCTCGGTGGTCGCCGTATCATT’. Then, trimmed and high quality of reads were aligned to mouse reference genome (GRCm39, Release-107 in (http://www.ensembl.org/Mus_musculus/Info/Index website) by gene transfer format (GTF) using by meRanGh align within meRanTK tool54 kits (Version 1.2.1b). This step also included splice-aware HISAT2 algorithm and arguments ‘-mmr 0.1’. Subsequently, SAM files were obtained and proceeded to conduct the identification of m5C sites using the arguments as follows: ‘-rl 150 -md 5 -ei 0.1 -cr 99 -fdr 0.01 -mcov 10 -mr 0.05’. The criteria used to filter m5C sites are: minimum 10 reads coverage, methylation ratio over 5% (0.05), and false discovery rate 1%. Then, gene annotation for m5C sites was retrieved by using meRanAnnotate with the general feature format 3 (GFF3).

### mRNA Bisulphite sequencing (remarks)

We used almost 40µg of total RNA for poly(A) purification and approximately 38 to 138 ng/ul of bs-treated mRNA for next generation sequencing. Most of the purified mRNAs in our experimental groups range between 1-4% of corresponding total RNAs, which is in line with previous mRNA-Bis-seq studies^56^. Prior to poly(A) purification and NGS, total RNA and mRNA integrity were assessed by Bioanalyzer to prove undegraded samples. Afterwards, poly(A) purified mRNAs were pooled with luciferase spike-in control (1:10000 ratio) derived from another species (*Photinus pyralis*, a.k.a. Firefly) and does not contain any m^5^C sites before bisulphite conversion. The average reads in our samples are 66.7M per sample. After trimming and quality control, the high-quality reads are mapped into in silico bs-converted mouse reference genome (GRCm39) and luciferase spike-in control using meRanTK tools^66^. We found that both our samples and spike-in controls were almost ∼99.8% converted (Supplementary TableS4), which ensures the efficiency of bisulphite conversion of non-methylated cytosines in the samples. This finding is convincingly similar to previous studies^56,67^.

Various studies on different tissue and cell types have reported a range of cut-off values^28,30,31,56,68^ using meRanTK tool^66^. This implies cell-context dependency in mRNA m^5^C methylation. Therefore, it is crucial to identify candidate m^5^C mRNA selection criteria in our samples. After a round of optimisation, we selected following criteria in meRanCall: minimum reads coverage (mcov):10, methylation ratio (mr)≥0.05, and false discovery rate (FDR)=1%. Using this selection, we detected nearly 100,000 m^5^C sites for each in vivo sample and 50,000 to 100,000 m^5^C sites in the in vitro samples. Looking into luciferase spike-in control, we generally determined 1-2 m^5^C sites per sample, except for wild-type-3 sample (Supplementary TableS3). Following these, we realised that some sites are missing from the analysis. This is because of obscured sites where m^5^C is completely lost in the *Nsun2*-loss samples. To revise this, we also included “mr:0 only” values. In this way, we did not discard the mr:0 sites but this might cause a background noise. To overcome this challenge, we also set the standard deviation between replicates within each group lower than 0.1.

### Meta-analysis of TCGA data

Human colorectal cancer datasets (TCGA-COAD and TCGA-READ PanCancer) were retrieved from cBioPortal. The expressions of gene of interests within TCGA-COAD and TCGA-READ patient datasets were analysed by using from cBioPortal and XenaBrowser platforms. Several CRC classifications were also obtained from Xenabrowser and Myant lab. For the survival analysis, Cox regression hazard model and Kaplan–Meier analysis were conducted using PanCancer datasets in KM-PLOT. Figures plotted in GraphPad Prism 9.5.1.

### Statistical analysis

Unpaired t-test and One-way ANOVA multiple comparison tests used by GraphPad Prism Software version 9.5.1 for Windows (GraphPad Software, San Diego, CA). Statistically significant changes were shown by a symbol (*). Symbol expressions are as follows: *P < 0.05 (significant); **P < 0.01 (very significant); ***P < 0.001 and ****P < 0.0001 (extremely significant), and ns: non-significant.

## Acknowledgements

This study is generously funded by Republic of Türkiye, Ministry of National Education Selection and Placement of Candidates Sent Abroad for Postgraduate Education (YLSY) scholarship program. This work was also funded by Cancer Research UK (CRUK) under Career Development Fellowship, A19166 (K.B.M), the European Research Council under Starting Grant, COLGENES – 715782 (K.B.M) and the MRC under Project Grant MR/X008762/1 (K.B.M.). We thank all the member of Myant lab for their support during the progression of this project. We are grateful to collaborate with Szu-Ying Chen, who received student scholarship from NSTC#A1102-U015 and NCKU-SP112002, and Professor Po-Hsien Huang in Taiwan.

## Author contributions

A.B.A. conceived the research and most the experiments. C.B. produced the virus required for knockdown experiments. Preliminary RNA-seq data obtained from A.H.. S.Y.C. analysed mRNA-Bisulphite sequencing. K.M., F.D., and P.H.H. provided critical advice. A.B.A and K.M. wrote the manuscript with contributions from all other authors.

## Data availability

RNAseq and mRNA bisulphite sequencing will be uploaded and made freely available prior to publication. All data reported in this paper and any additional information will be shared upon reasonable request to lead contact.

---

**Supplementary Fig S1.**
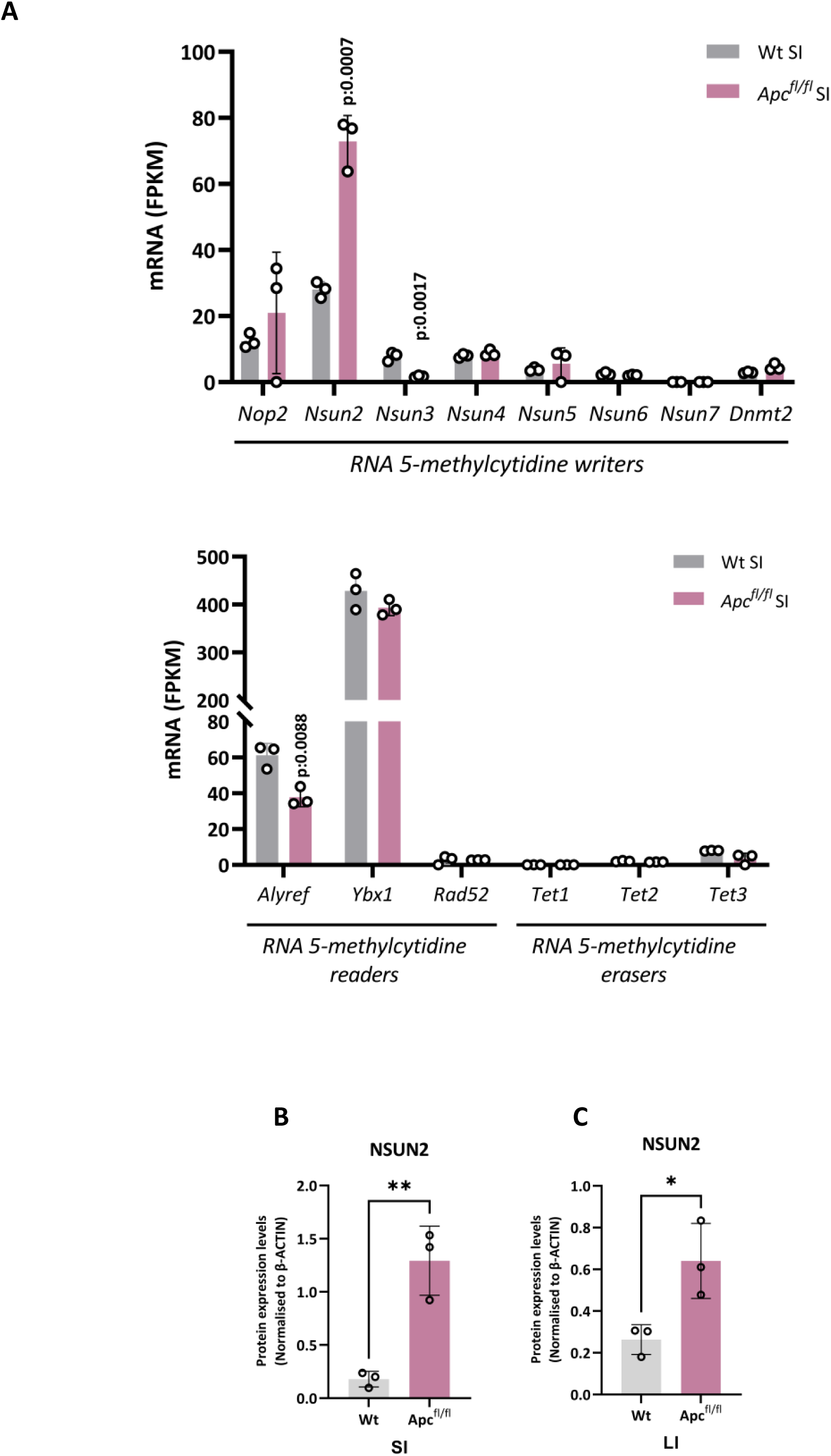
RNA metabolic processes are enriched following *Apc*-loss and NSUN2 is upregulated in *Apc*-deficient mouse intestinal tissues. **A** mRNA expression levels (FPKM values) of genes involved in m5C methylation (writers, readers and erasers) in RNA-seq from wild-type and *Apc^fl/fl^* mouse small intestines collected post day 5 Tamoxifen induction (data are presented as mean ± SD; ***p:0.0007, p:0.0017; two-tailed t-test, n = 3vs3 biologically independent mice). **B** Quantification of normalised protein expression levels in wild-type and *Apc^fl/fl^* mouse small intestine, using β-ACTIN loading control (data are presented as mean ± SD; **p:0.0044; two-tailed t-test, n = 3vs3 biologically independent mice). **C** Quantification of normalised protein expression levels in wild-type and *Apc^fl/fl^* mouse colon, using β-ACTIN loading control (data are presented as mean ± SD; *p:0.0279; two-tailed t-test, n = 3vs3 biologically independent mice).

**Supplementary Fig S2.**
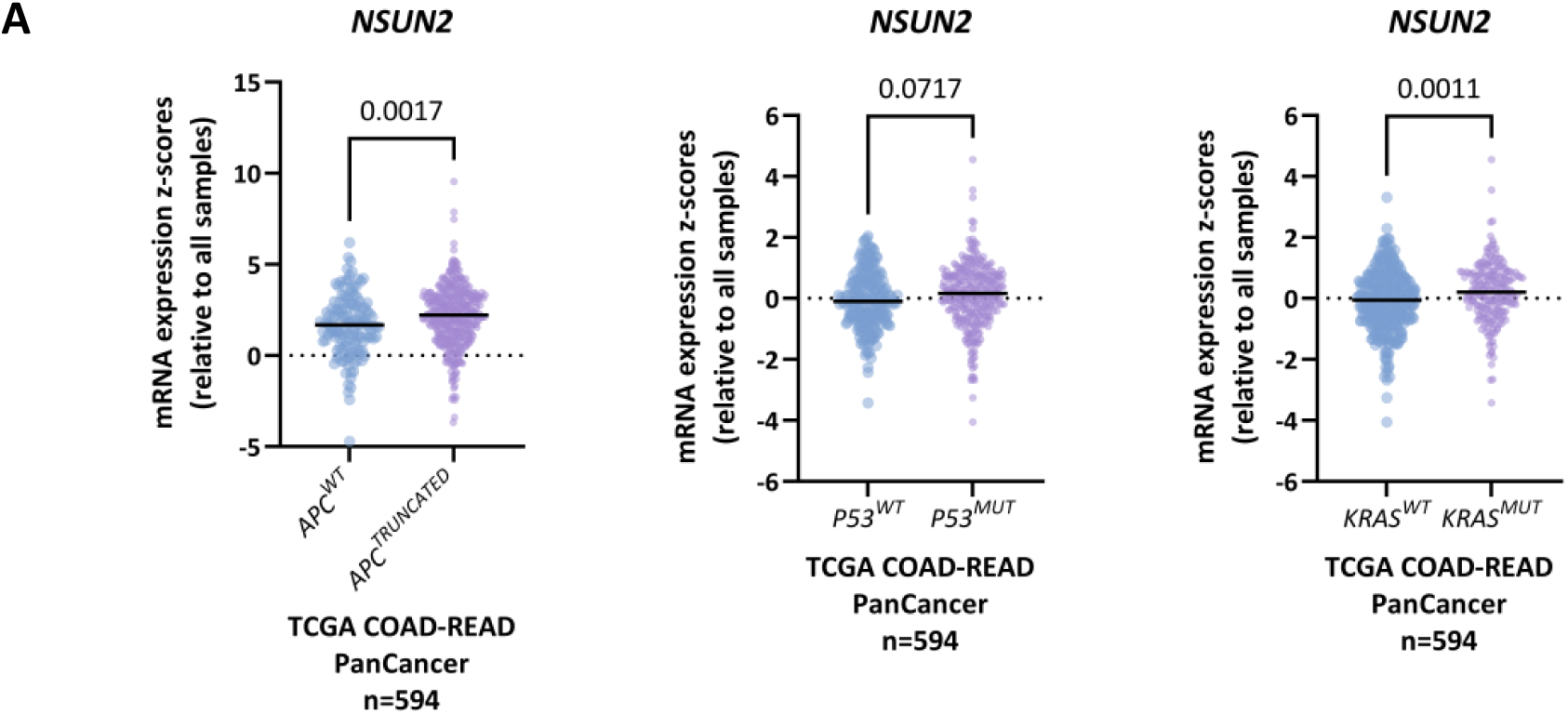
NSUN2 is overexpressed in patients with truncated *APC* mutation but not in *KRAS* and *P53* mutation. **A** *NSUN2* mRNA expression in primary tumours with mutations in *APC (*truncated*) (left)*, *KRAS (middle)*, and *P53 (right)* genes from TCGA COAD-READ PanCancer patients compared to *APC^wt^* primary tumours in cBioportal. (data are presented as mean ± SD; **p:0.0017, *p:0.0279; two-tailed t-test, n *APC^wt^* vs *APC^Truncated^* n=141vs385 patients, *KRAS^wt^* vs *KRAS^mut^* n=310vs212, *P53^wt^*vs *P53^mut^* n=213vs309).

**Supplementary Fig S3.**
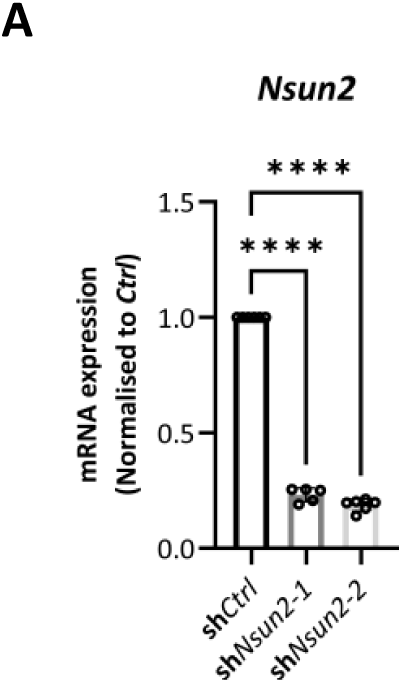
*Nsun2* knockdown in *Apc^fl/fl^* mouse small intestinal organoids validated by qPCR. **A** Validation of *Nsun2* knockdown in *Apc^fl/fl^* mouse small intestinal organoids by qPCR (left) (data are presented as mean ± SD; ****p<0.0001) and by western blot (middle, right) (data are presented as mean ± SD; ****p<0.0001).

**Supplementary Fig S4.**
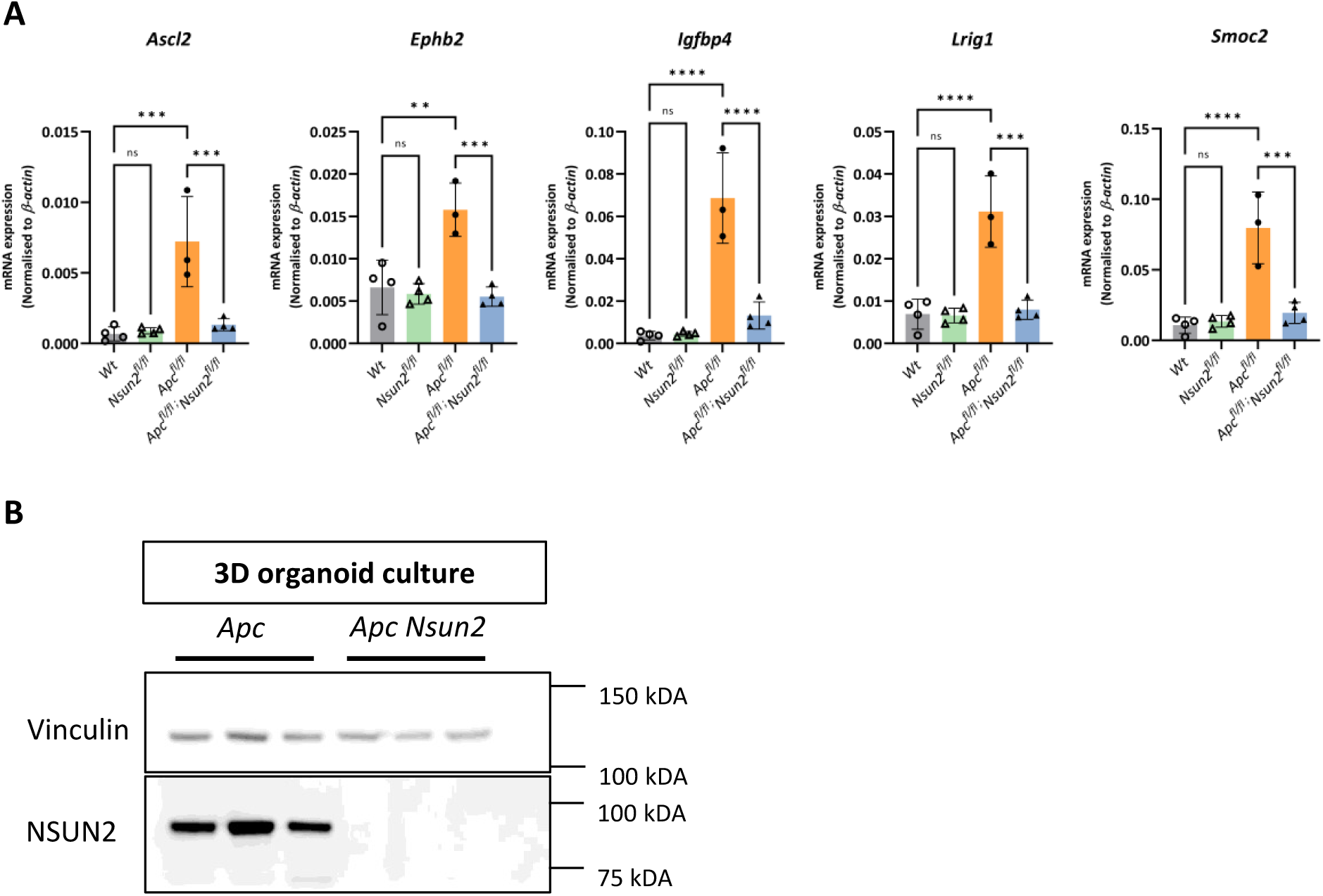
Reduction in other stem cell marker genes after *Nsun2*-deletion and validation of NSUN2 deletion in intestinal crypt culture. **A** mRNA expression levels of other stem cell marker genes, *Ascl2, Ephb2, Igfbp4, Lrig1,* and *Smoc2*. All mRNA expressions were normalised to *β-actin*, a housekeeping control. (data are presented as mean ± SD; two-tailed t-test, ***p<0.0001, **p<0.001 , *p<0.05, not significant:ns, (n = 4vs4vs3vs4 biologically independent mice). **B** Images of western blotting analysis for NSUN2 and a house-keeping control VINCULIN protein expressions in *Apc^fl/fl^*and *Apc^fl/fl^ Nsun2^fl/fl^*Vil-CreERT2 mouse small intestinal crypt cultures.

**Supplementary Fig S5.**
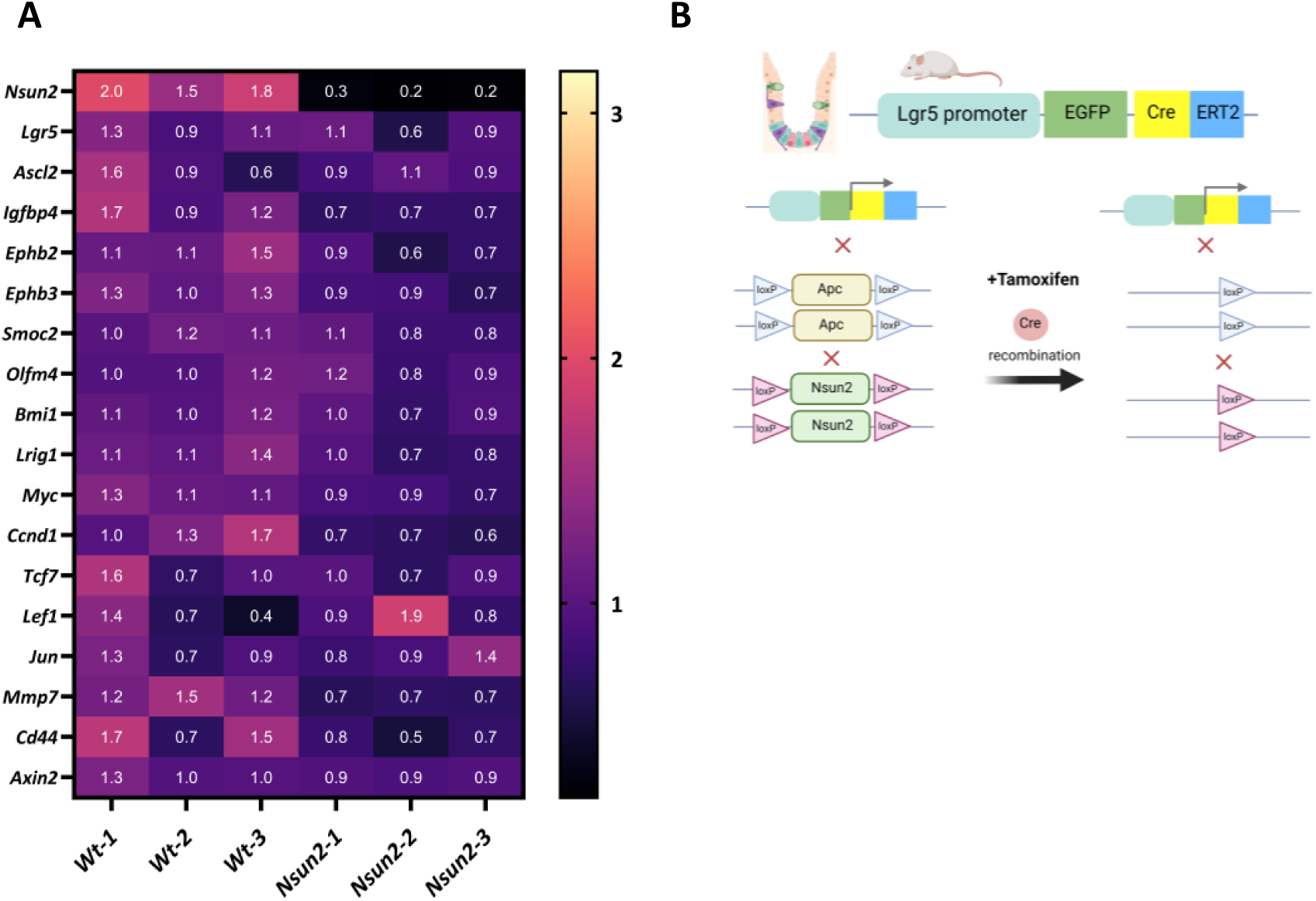
*Nsun2*-loss do not affect transcriptome of normal intestine. **A** Heat-map of differentially expressed intestinal stem cell markers and Wnt targets comparing small intestinal crypts from *Vil-Wt* vs *Vil-Nsun2* mice (n = 3 vs 3 biologically independent mice). **B** Schematic demonstration of *Apc* and *Nsun2* knockout mouse model generation in the intestinal tissue-specific *Lgr5-IRES-eGFP-CreERT2* line. The figure was created by Biorender.com.

**Supplementary Fig S6.**
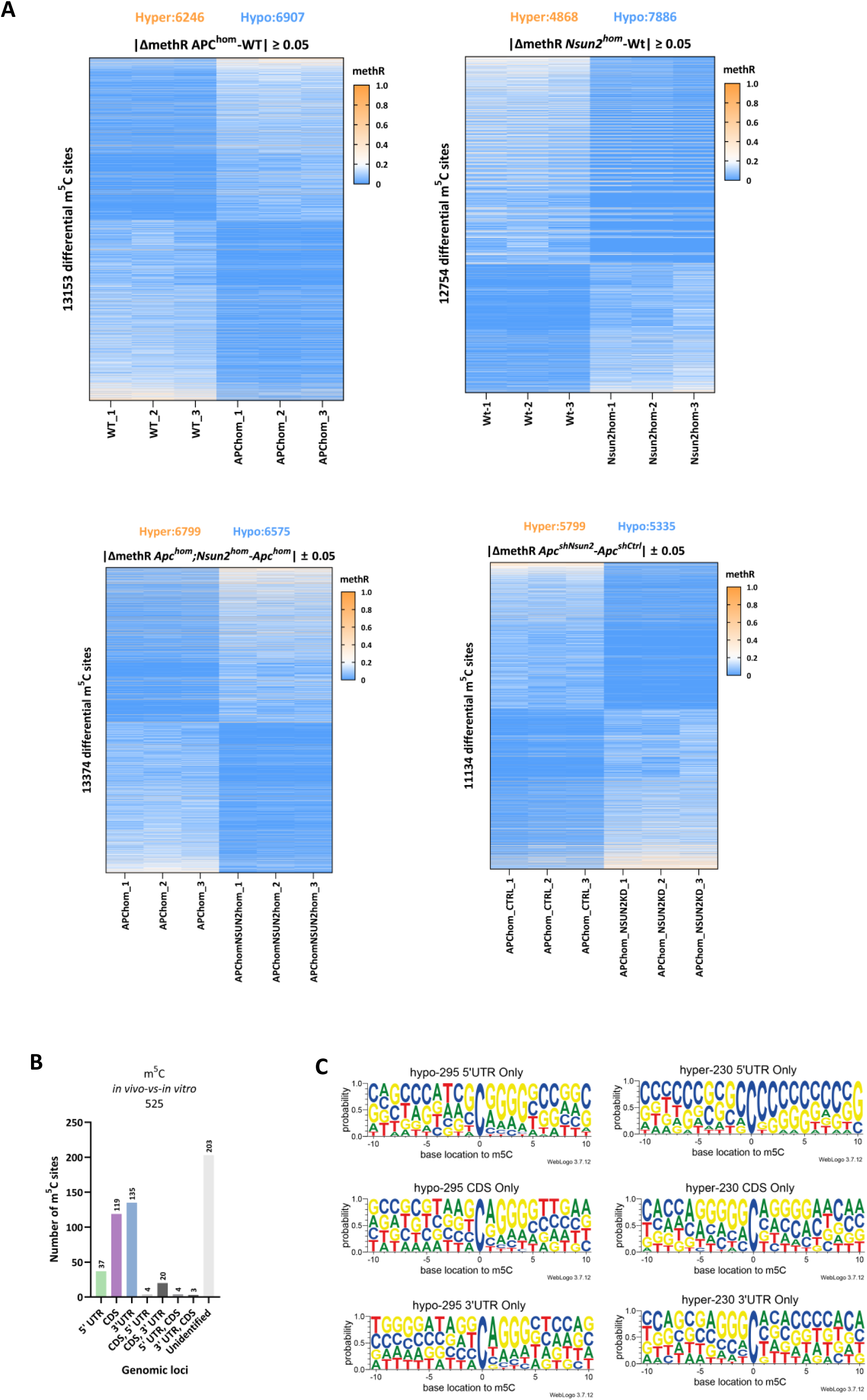
mRNA methylation profile in all experimental groups and distribution and sequence context of 525 common m^5^C sites between in vivo and in vitro *ApcNsun2* deficient models. **A** Heat-map of methylation profile in all experimental models used in whole transcriptome bisulphite sequencing. 13,153 m^5^C sites for *Vil-Wt* vs *Vil-Apc* mice, 12,754 m^5^C sites for *Vil-Wt* vs *Vil-Nsun2* mice, 13,374 m^5^C sites for *Vil-Apc* vs *Vil-ApcNsun2* mice are identified (n = 3 vs 3 biologically independent mice). 11,134 m^5^C sites for *Apc^fl/fl^-shCtrl* and *Apc^fl/fl^-shNsun2* are identified (n = 3 experimental replicates). Differentially methylated sites (DMSs) were defined using Δmethylation ration: ±0.05. **B** The bar graph represents distribution of 525 common m^5^C sites detected as a result of overlapping vivo and vitro m^5^C sites in different mRNA regions **C** The probability pattern represents sequence context of 10 bases up and downstream of hypo-methylated (295) and hyper-methylated (235) of common 525 NSUN2-dependent m^5^C sites based on located in different mRNA regions

**Supplementary Fig S7.**
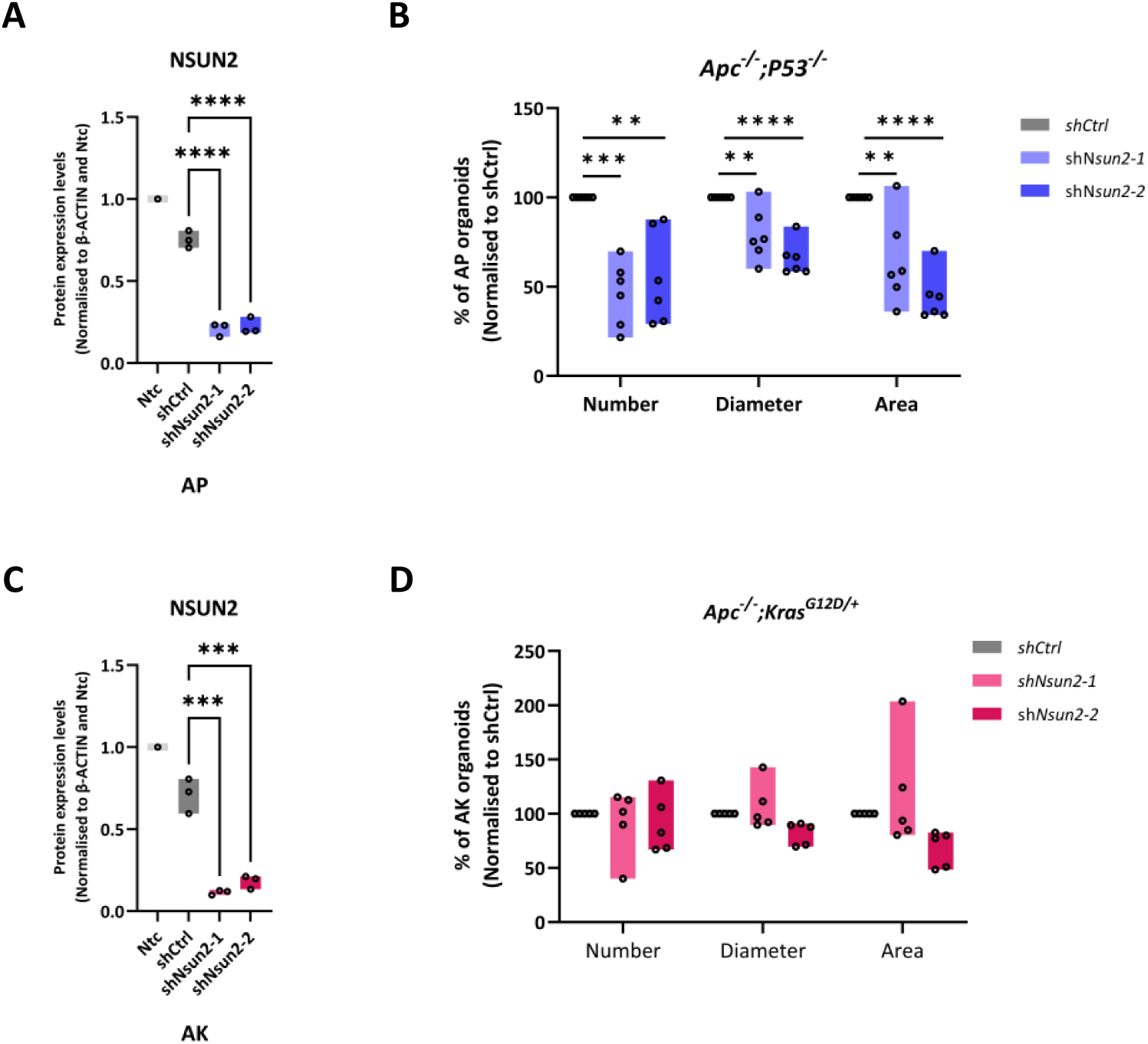
NSUN2 knockdown reduces clonogenic capacity of *Apc^fl/fl^ ; P53^fl/fl^* but not *Apc^fl/fl^ ; Kras^G12D/+^* mouse colorectal cancer organoids. **A** Quantification of normalised NSUN2 protein expression levels in *Apc^fl/fl^ ; P53^fl/fl^ (AP)-shCtrl* and *Apc^fl/fl^ ; P53^fl/fl^ (AP)-shNsun2* mouse small intestinal organoids by western blot, using β-ACTIN loading control (data are presented as mean ± SD; ****p<0.0001 (*shCtrl* and *shNsun2-1*), ****p<0.0001 (*shCtrl* and *shNsun2-2*); One-way ANOVA test comparing to *shCtrl*, n = 3 experimental replicates). **B** The percentage of organoid number, diameter, and area in *shNsun2-1* and *shNsun2-2* groups compared to *shCtrl* in *Apc^fl/fl^ ; P53^fl/fl^ (AP)* mouse small intestinal organoids (data are presented as mean ± SD; ***p<0.0001, **p<0.001 , *p<0.05, Ordinary One-way ANOVA test compared to *shCtrl,* n = 6 experimental replicates). **C** Quantification of normalised NSUN2 protein expression levels in *Apc^fl/fl^ ; Kras^G12D/+^ (AK)-shCtrl* and *Apc^fl/fl^ ; Kras^G12D/+^ (AK)-shNsun2* mouse small intestinal organoids by western blot, using β-ACTIN loading control (data are presented as mean ± SD; ***p:0.0001 (*shCtrl* and *shNsun2-1*), ***p:0.0002 (*shCtrl* and *shNsun2-2*); One-way ANOVA test comparing to *shCtrl*, n = 3 experimental replicates). **D** The percentage of organoid number, diameter, and area in *shNsun2-1* and *shNsun2-2* groups compared to *shCtrl* in *Apc^fl/fl^ ; Kras^G12D/+^ (AK)* mouse small intestinal organoids (data are presented as mean ± SD; ***p<0.0001, **p<0.001 , *p<0.05, not significant values are not indicated, Ordinary One-way ANOVA test compared to *shCtrl,* n = 5 experimental replicates).

